# Multiple non-redundant functions underlie the essentiality of lipopolysaccharide in Gram-negative bacteria

**DOI:** 10.64898/2026.10.21.760538

**Authors:** Ryan Adam Valdez, Nina Zheng, Sarah D. Beagle, Jiawei Sun, Kerwyn Casey Huang, Petra Anne Levin

**Affiliations:** Department of Biology, Washington University in St. Louis, St. Louis, MO, 63130, USA; Department of Bioengineering, Stanford University, Stanford, CA 94305, USA; Department of Microbiology and Immunology, Stanford University School of Medicine, Stanford, CA 94305, USA; Chan Zuckerberg Biohub, San Francisco, CA 94158, USA

## Abstract

Lipopolysaccharide (LPS) is well known for its immunostimulatory activity and its contribution to the permeability barrier of the Gram-negative outer membrane. Yet LPS is essential for growth in nearly all Gram-negative bacteria, even under laboratory conditions where its barrier function should be less critical. LPS contributes to envelope mechanics, membrane biogenesis, and cell division, but whether any of these functions accounts for its essentiality remains unclear. Here, we systematically tested these possibilities by depleting LPS in *Escherichia coli* and *Klebsiella pneumoniae*. Osmoprotection largely preserved cell-envelope integrity but not viability, demonstrating that the mechanical contribution of LPS alone cannot explain its essentiality. Genetic perturbations that increase peptidoglycan rigidity or alter phospholipid homeostasis partially improved growth, supporting independent roles for LPS in envelope mechanics and membrane biogenesis. An unbiased suppressor screen further identified activation of the MarA regulon as a potent mechanism for surviving LPS depletion. Protection required transport by the AcrAB-TolC multidrug efflux pump, revealing an unexpected connection between efflux and cell-envelope homeostasis. Thus, LPS essentiality emerges from multiple non-redundant contributions to Gram-negative cell-envelope physiology.

## Introduction

The Gram-negative cell envelope is a multilayered structure composed of a phospholipid-rich inner membrane, a peptidoglycan cell wall within the aqueous intermembrane space known as the periplasm, and an asymmetric outer membrane (OM) containing phospholipids in the inner leaflet^1^ and lipopolysaccharide (LPS) in the outer leaflet^2^. This lipid asymmetry is critical for the OM’s function as a permeability barrier^3^ and is actively maintained by systems that remove phospholipids from the outer leaflet, including the OM phospholipase PldA^4,5^ and the Maintenance of Lipid Asymmetry (Mla) pathway^6^. Synthesized in the cytoplasm and transported to the cell surface^2^, LPS occupies an estimated ∼30% of the cell surface of *Escherichia coli*^7^. LPS is both a major target of last-resort antibiotics^8^, including colistin^9^, and a potent activator of innate immune responses across diverse eukaryotes^10^. Yet despite its importance in interactions with the environment and host, LPS is essential for growth in nearly every Gram-negative bacterium studied to date^7^, and the basis of this essentiality remains unclear.

In *E. coli*, disruption of LPS biosynthesis results in aberrant cell division, growth arrest, and eventual cell lysis^11–13^. Only a small number of Gram-negative bacteria are known to survive without LPS^7^, including the pathogen *Acinetobacter baumannii* in which multiple compensatory mutations and physiological adaptations enable survival^14–17^. These observations establish that LPS is generally indispensable but do not explain why depletion of a single component of the OM is lethal. In laboratory conditions, where cells are not challenged with host defenses or environmental toxins, loss of the permeability barrier alone is unlikely to account for LPS essentiality. Instead, accumulating evidence implicates LPS in multiple aspects of cell-envelope biogenesis and function that could independently or collectively explain its essentiality.

One possibility is that LPS is essential because of its contribution to cell-envelope mechanics. Biophysical and cytological studies indicate that the OM functions together with the thin Gram-negative cell wall as a load-bearing structure that resists turgor pressure^18–20^. OM rigidity arises in part through electrostatic interactions between divalent cations and negatively charged phosphate groups on lipid A^18^, the anchoring portion of LPS, and mechanical measurements suggest that the OM can contribute stiffness comparable to that of the cell wall^18^. Consistent with this model, defects in LPS biosynthesis frequently result in cell lysis^13^, and interactions between LPS and magnesium ions are required for proliferation of cell wall-deficient *E. coli* L-forms^21,22^. Moreover, mutations that increase peptidoglycan crosslinking improve growth of LPS-deficient *Acinetobacter baumannii*^15,23,24^. These findings raise the possibility that the mechanical contribution of LPS is itself responsible for its essentiality.

A second possibility is that LPS is required for coordinated expansion of the cell envelope. Because LPS occupies a large fraction of the outer leaflet^25^, restricting its synthesis should limit OM expansion while phospholipid synthesis and growth of the rest of the envelope continue. Consistent with this model, LPS-deficient *A. baumannii* frequently acquires mutations in *pldA* and components of the Mla pathway that increase phospholipid retention in the outer leaflet and thus contribute to preserving its area^14,26^. Altering lipid synthesis can also compensate for reduced LPS abundance: mutations affecting lipid synthesis and transport promote growth of LPS-deficient *A. baumannii*^16^, whereas mutations that reduce fatty acid synthesis confer resistance to inhibitors of the essential LPS biosynthetic enzyme LpxC in both *E. coli* and *Klebsiella pneumoniae*^27,28^. Likewise, growth in minimal medium improves proliferation of LPS-deficient *A. baumannii,* potentially by reducing phospholipid production and allowing more balanced membrane expansion^14^. Together, these observations suggest that LPS may be essential as a major structural component required to coordinate growth of the OM with the remainder of the cell envelope.

A third possibility is that LPS synthesis is directly integrated with other essential cell-envelope processes, including cell division. In *Pseudomonas aeruginosa*, LpxC interacts with MurA, which catalyzes the first committed step of peptidoglycan synthesis, and MurA stimulates LpxC activity *in vitro*, suggesting coordination between synthesis of these two major envelope components^29^. Although this interaction is not known to occur in *E. coli*, disruption of multiple steps in LPS biosynthesis results in defective assembly of the division machinery and filamentous growth^12,30,31^. Whether these phenotypes arise secondarily from altered membrane mechanics or imbalanced envelope biogenesis or instead reflect more direct coordination between LPS synthesis and cytokinesis remains unclear.

Thus, although LPS has been implicated in multiple potentially essential functions, whether any one of these functions explains its essentiality or whether essentiality instead emerges from their combined contributions remains unresolved. Here, we use complementary cytological, physiological, and genetic approaches to systematically test these possibilities in *E. coli* MG1655 and *K. pneumoniae* TOP52. We find that the mechanical contribution of LPS is critical for maintaining cell-envelope integrity but is insufficient to sustain viability, while perturbations affecting peptidoglycan rigidity and phospholipid homeostasis partially compensate for LPS depletion. An unbiased suppressor screen further identified the MarA–AcrAB-TolC pathway as an unexpected determinant of survival during LPS depletion. Together, our findings argue that LPS essentiality does not arise from a single indispensable activity but instead emerges from multiple non-redundant contributions to Gram-negative cell envelope physiology.

## Results

### LPS depletion results in progressive growth arrest and loss of envelope integrity

To determine how LPS depletion affects bacterial growth and cell-envelope integrity, we established two complementary methods to conditionally inhibit LPS synthesis. First, we used an arabinose-inducible tunable CRISPR interference (tCRISPRi) system in *E. coli*^32^ to inhibit expression of *lpxC*, which encodes the enzyme catalyzing the first committed step in LPS synthesis. Second, we used the LpxC inhibitor CHIR-090^33^ to inhibit LPS synthesis in *E. coli* MG1655 and *K. pneumoniae* TOP52. Because LPS is abundant and relatively stable in the OM, inhibiting its synthesis should result in progressive depletion as existing LPS is diluted during growth. We therefore quantified LPS abundance immediately before treatment and 180 min later (Fig 1A).

**Figure 1:**
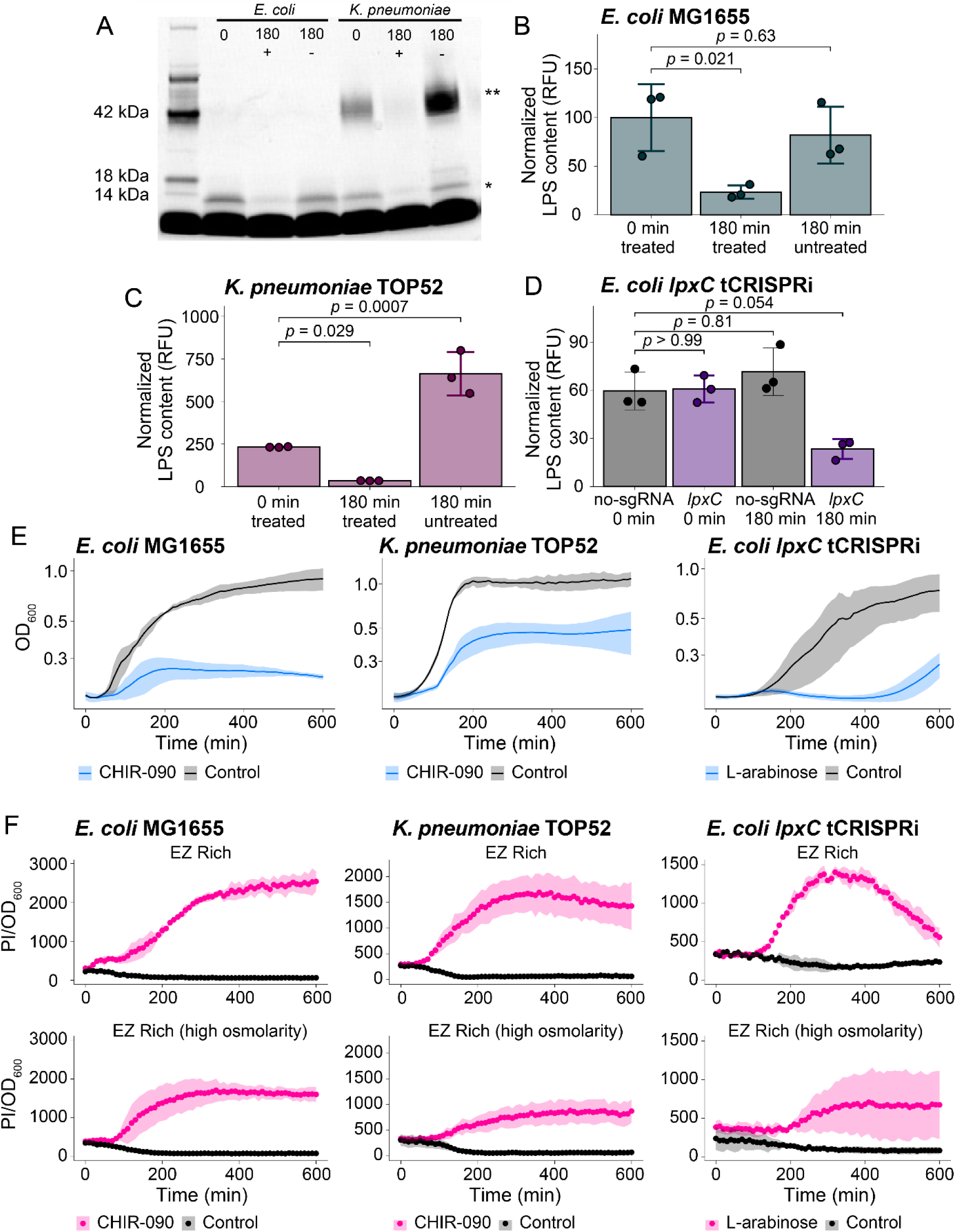
LPS depletion results in growth arrest and loss of cell-envelope integrity. **(A)** Representative Pro-Q Emerald-stained gel used to quantify LPS abundance. Single and double asterisks indicate the migration positions of LPS lacking O antigen (*; *E. coli* MG1655) and LPS containing O antigen (**; *K. pneumoniae* TOP52), respectively. Samples were collected before treatment (0 min) or after 180 min in the presence (+) or absence (–) of CHIR-090. **(B-D)** Relative LPS abundance in *E. coli* MG1655 (B)*, K. pneumoniae* TOP52 (C), and the *lpxC* tCRISPRi strain (D). Both CHIR-090 treatment and *lpxC* repression substantially reduced LPS abundance. LPS signal was normalized to the 42-kDa glycoprotein ladder band and total protein and scaled such that the mean pretreatment value for *E. coli* MG1655 was 100. Points represent independent experimental replicates; bars and error bars indicate mean ± standard deviation (SD). *p-*values were determined using Dunnett’s multiple-comparisons test. **(E)** Growth during LPS depletion induced by CHIR-090 in *E. coli* MG1655 and *K. pneumoniae* TOP52 or by *lpxC* tCRISPRi induction with L-arabinose. LPS depletion resulted in progressive growth arrest with both pharmacological and genetic inhibition of LPS synthesis. Lines and shaded regions indicate mean ± SD from at least three biological replicates. **(F)** PI fluorescence normalized to OD_600_ during LPS depletion under standard-osmolarity conditions (EZ Rich) or in medium supplemented with 500 mM D-sorbitol (EZ Rich HOSM). Increased PI/OD_600_ during LPS depletion reflects increased PI uptake and progressive loss of cell-envelope integrity, which was reduced under high-osmolarity conditions. Lines and shaded regions indicate mean ± SD from at least three biological replicates.

Both approaches substantially reduced cellular LPS over the course of the experiment (Fig 1B-D). CHIR-090 treatment reduced LPS abundance by ∼80% relative to starting levels in both *E. coli* and *K. pneumoniae* after 180 min (Fig 1B,C). LPS abundance remained essentially unchanged in untreated *E. coli* but increased ∼2.8-fold in untreated *K. pneumoniae*, resulting in ∼95% less LPS in CHIR-090-treated than untreated *K. pneumoniae* after 180 min. Similarly, induction of *lpxC* tCRISPRi reduced LPS abundance in *E. coli* by ∼66% relative to both the starting culture and the non-targeting control (Fig 1D).

Progressive LPS depletion was accompanied by growth arrest. In both *E. coli* and *K. pneumoniae*, CHIR-090 treatment caused an immediate reduction in growth rate followed by a plateau in culture density relative to untreated controls (Fig 1E). Similarly, induction of *lpxC* tCRISPRi arrested growth for approximately six hours, after which growth resumed at a reduced rate (Fig 1E). The basis of this recovery is unknown, although the growth dynamics suggest a mutation in the CRISPRi system.

Growth arrest was accompanied by progressive loss of cell-envelope integrity. We monitored cell envelope integrity using propidium iodide (PI), a membrane-impermeant nucleic acid stain that enters cells with compromised envelopes^34^. Untreated cells exhibited little PI fluorescence, whereas LPS-depleted cells progressively accumulated PI (Fig 1F). During CHIR-090 treatment, PI fluorescence normalized to culture turbidity peaked near the onset of growth arrest, consistent with envelope damage occurring predominantly during active growth. Following *lpxC* tCRISPRi induction, PI fluorescence continued to increase during growth arrest but declined as growth resumed, consistent with recovery of a subpopulation that is resistant to tCRISPRi induction.

Thus, complementary chemical and genetic approaches demonstrate that progressive LPS depletion leads to growth arrest accompanied by loss of cell-envelope integrity.

### High osmolarity preserves cell-envelope integrity but not culturability during LPS depletion

If the primary essential function of LPS is to provide mechanical rigidity to the cell envelope^18^, reducing mechanical stress should protect cells during LPS depletion. Previous work showed that increasing medium osmolarity supports growth of cell wall-deficient *E. coli* L-forms by reducing turgor pressure^21^. We therefore asked whether osmoprotection similarly compensates for LPS depletion.

We monitored individual *E. coli* and *K. pneumoniae* cells during LPS depletion on agarose pads supplemented with 500 mM D-sorbitol (hereafter referred to as sorbitol), matching the osmolyte concentration previously shown to support growth of *E. coli* L-forms^21^. Time-lapse microscopy in the presence of PI allowed us to simultaneously monitor proliferation and cell-envelope integrity (Fig 2A).

**Figure 2:**
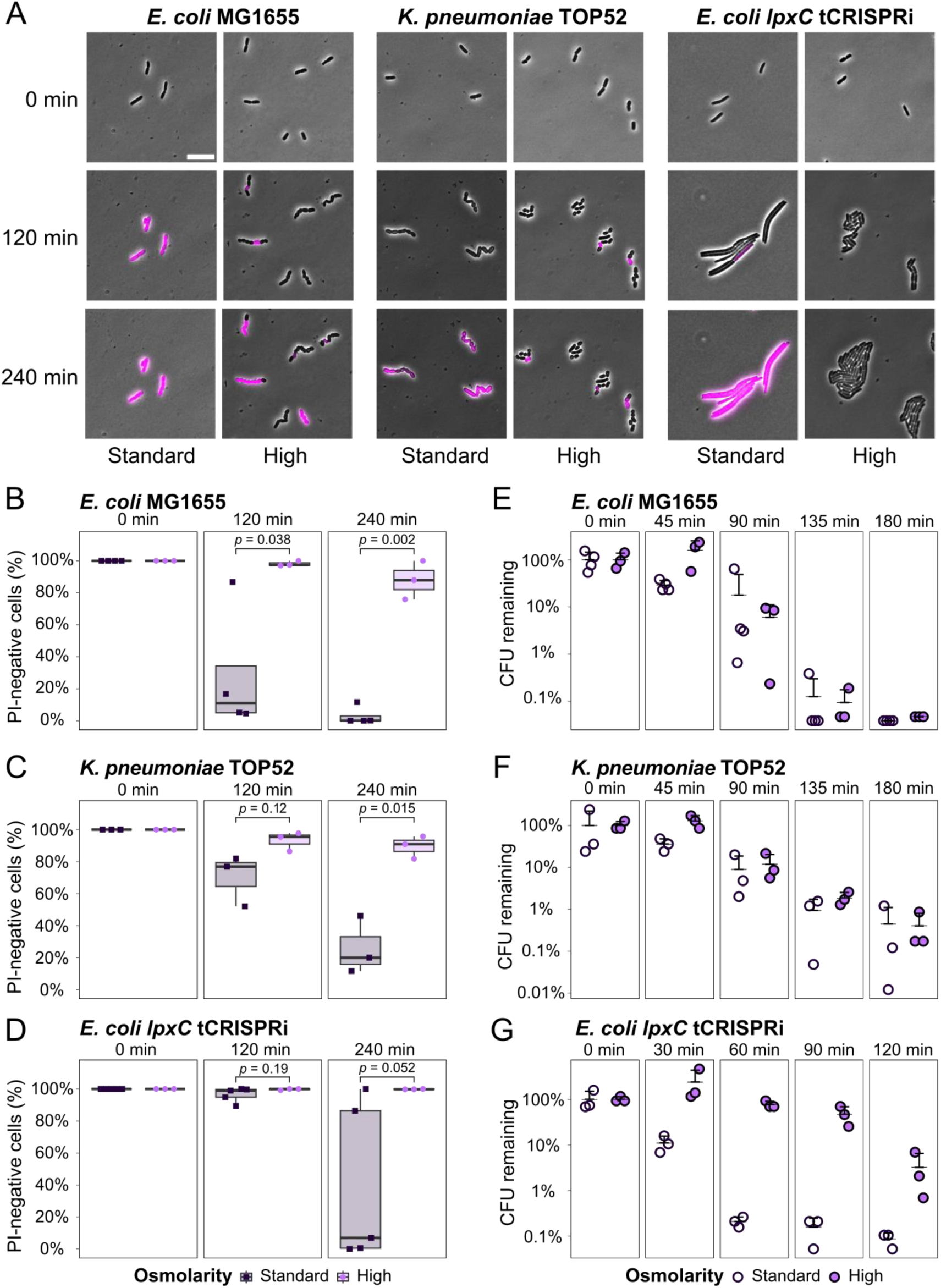
High osmolarity preserves cell-envelope integrity but not culturability during LPS depletion. **(A)** Representative microscopy images of *E. coli* MG1655, *K. pneumoniae* TOP52, and the *lpxC* tCRISPRi strain during LPS depletion under standard- and high-osmolarity conditions. PI staining is shown in magenta. Scale bar, 10 µm. **(B-D)** Percentage of PI-negative cells during LPS depletion in *E. coli* MG1655 (B), *K. pneumoniae* TOP52 (C), and the *lpxC* tCRISPRi strain (D) under standard- and high-osmolarity conditions on solid medium. High osmolarity substantially increased the fraction of cells that remained PI-negative during LPS depletion, indicating preservation of cell-envelope integrity. Each point represents a biological replicate. Box plots show the median, interquartile range (IQR), and values within 1.5X the IQR. *p*-values were determined using Welch’s unequal-variances *t*-test. **(E-G)** Culturability during LPS depletion in liquid medium for *E. coli* MG1655 (E), *K. pneumoniae* TOP52 (F), and the *lpxC* tCRISPRi strain (G) under standard- and high-osmolarity conditions. Despite preserving cell-envelope integrity, high osmolarity did not prevent the progressive loss of culturability during LPS depletion. CFU/mL at each time point is shown relative to the corresponding *t*_0_ values. Each point represents a biological replicate. Wider black bars indicate the mean; upper error bars indicate mean + SD. Lower error bars are omitted because they extend below zero and therefore cannot be displayed on the logarithmic scale.

High osmolarity modestly increased proliferation during LPS depletion but did not restore sustained growth. CHIR-090-treated *E. coli* and *K. pneumoniae* divided only one to two times regardless of medium osmolarity. CHIR-090 treatment typically produced short chains of ∼4–8 cells, whereas *lpxC* tCRISPRi induction typically produced short filaments at standard osmolarity and chains of cells at high osmolarity (Fig 2A, S1A). Notably, induced *lpxC* tCRISPRi cells underwent approximately four generations at high osmolarity compared with two generations at standard osmolarity. Untreated cells and non-targeting controls formed microcolonies of normal rod-shaped cells (Fig S1B).

In contrast to its modest effect on proliferation, high osmolarity dramatically preserved cell envelope integrity during LPS depletion. After 240 min of CHIR-090 treatment, only 3% of *E. coli* cells remained PI-negative under standard conditions compared with 88% at high osmolarity (Fig 2B). Similarly, 26% and 89% of *K. pneumoniae* cells remained PI-negative at standard and high osmolarity, respectively (Fig 2C). Osmoprotection also substantially increased the fraction of PI-negative cells following *lpxC* tCRISPRi induction (Fig 2D). In liquid cultures, PI fluorescence likewise remained lower at high than at standard osmolarity during LPS depletion (Fig 1F). Thus, osmoprotection largely prevents the loss of cell-envelope integrity caused by LPS depletion.

We next asked whether preserving envelope integrity was sufficient to preserve culturability. Colony-forming units (CFUs) declined rapidly following CHIR-090 treatment in both *E. coli* and *K. pneumoniae* regardless of medium osmolarity, decreasing by more than two orders of magnitude within 180 min (Fig 2E,F). High osmolarity modestly slowed the decline in CFUs following *lpxC* tCRISPRi induction, but culturability nevertheless decreased by more than an order of magnitude over the course of the experiment (Fig 2G). High osmolarity did not interfere with LPS depletion (Fig S2A-C). Thus, osmoprotection largely uncouples loss of cell-envelope integrity from loss of culturability, demonstrating that the mechanical contribution of LPS alone is insufficient to explain its essentiality.

### Perturbations to phospholipid homeostasis enhance growth during LPS depletion

LPS may also be required for balanced expansion of the cell envelope. Because LPS occupies a large fraction of the outer leaflet of the OM, limiting its synthesis should restrict OM expansion while phospholipid synthesis continues. Consistent with this model, membrane biogenesis can limit growth of LPS-deficient *A. baumannii*^14^. We therefore predicted that perturbations that rebalance membrane biogenesis, either by increasing phospholipid retention in the OM, reducing the rate of IM expansion, or otherwise altering phospholipid synthesis, should improve growth during LPS depletion.

We screened *E. coli* mutants defective in genes implicated in membrane biogenesis using a Kirby-Bauer disk diffusion assay^35^ with CHIR-090. In this assay, smaller zones of inhibition (ZOIs) indicate improved growth in the presence of CHIR-090, allowing detection of mutations that provide even modest protection (Fig 3A). Consistent with our CFU experiments, high osmolarity did not alter the CHIR-090 ZOI of wild-type cells (Fig S2D), further indicating that osmoprotection alone is insufficient to improve growth during LPS depletion. Mutants that reduced the CHIR-090 ZOI by at least 10% were selected for further characterization of LPS abundance and culturability during CHIR-090 treatment.

**Figure 3:**
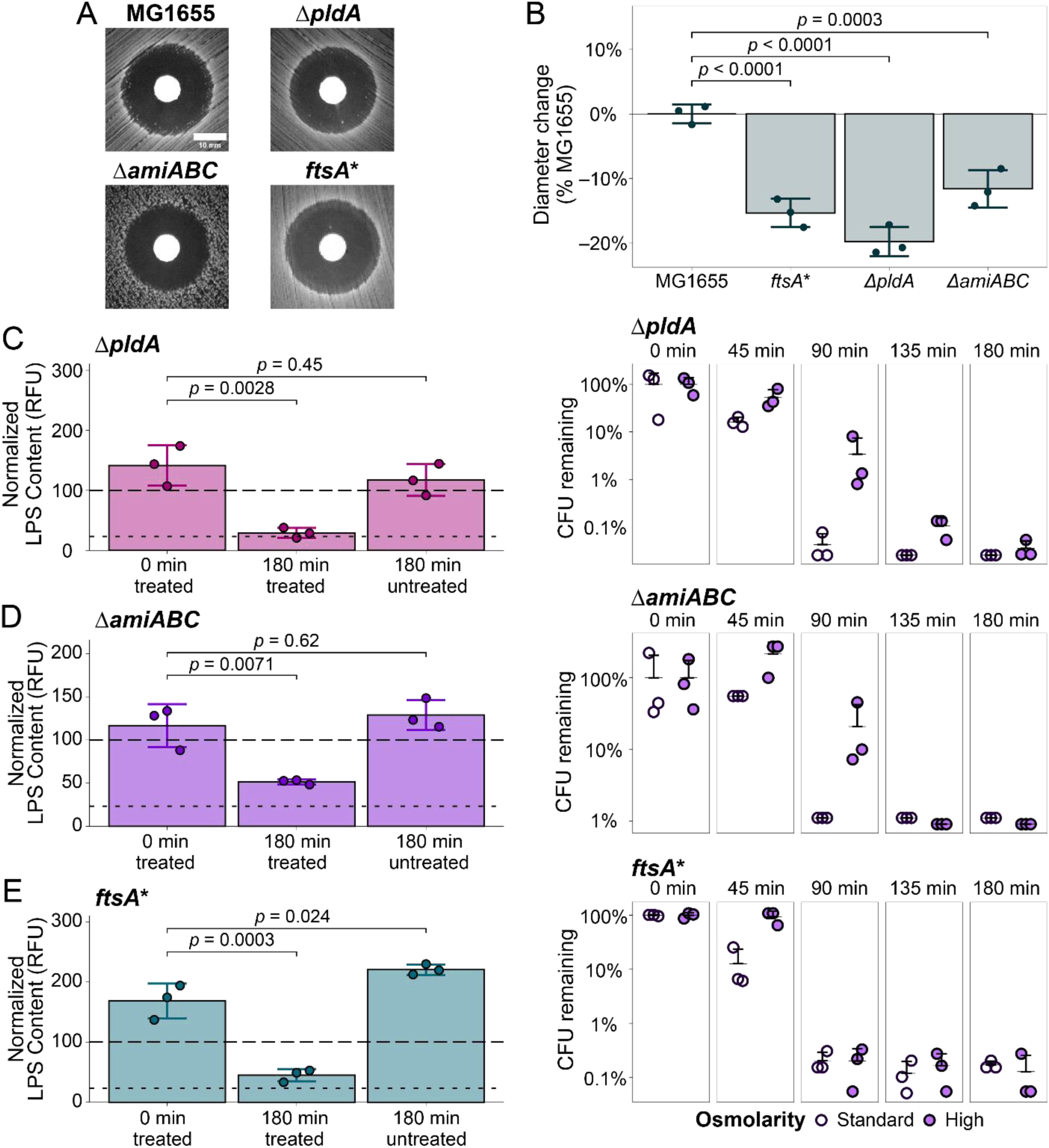
Mutations affecting distinct cell-envelope processes reduce susceptibility to LPS depletion. **(A)** Representative ZOIs produced by CHIR-090 in wild-type *E. coli* MG1655, Δ*pldA*, Δ*amiABC,* and *ftsA\**. Scale bar, 10 mm. **(B)** Change in ZOI diameter relative to MG1655. Negative values indicate smaller ZOIs. All three mutants exhibited reduced ZOIs, indicating increased growth in the presence of CHIR-090. Each point represents a biological replicate; bars and error bars represent mean ± SD. *p*-values were determined using Dunnett’s multiple-comparisons test. **(C-E)** Relative LPS abundance during CHIR-090 treatment in Δ*pldA* (C), Δ*amiABC* (D), and *ftsA\** (E), quantified as in Fig 1B-D. Δ*pldA* lost culturability despite its reduced ZOI, whereas Δ*amiABC* and *ftsA\** showed greater preservation of culturability during LPS depletion. High osmolarity further increased culturability in Δ*pldA* and Δ*amiABC*. Dashed and dotted lines indicate the mean pretreatment and 180-min CHIR-090-treated values, respectively, for MG1655. Corresponding plots at right show culturability during CHIR-090 treatment under standard- and high-osmolarity conditions, quantified as in Fig 2E-G. CFU/mL at each time point is shown relative to the corresponding *t*_0_ value.

Mutations predicted to increase phospholipid retention in the OM improved growth in the presence of CHIR-090. Consistent with previous observations in *A. baumannii*^14,26^, deletion of *pldA* (*pldA*::*kan*) or *mlaA* (*mlaA*::*kan*) reduced the CHIR-090 ZOI by 19.7% and 9.6%, respectively, relative to the parental strain (Fig 3B, Table S4). We further characterized Δ*pldA*, which exhibited the stronger phenotype. CHIR-090 depleted LPS to a similar extent in Δ*pldA* and wild-type cells (Fig 3C), indicating that its reduced susceptibility was not due to preservation of LPS abundance. Despite improved growth in the disk diffusion assay, however, Δ*pldA* lost culturability more rapidly than wild-type cells during CHIR-090 treatment in liquid medium (Fig 3C versus Fig 2E). High osmolarity substantially slowed this decline in *ΔpldA* (Fig 3C), suggesting that phospholipid retention in the OM and reduced mechanical stress have distinct, context-dependent effects during LPS depletion.

We next asked whether altering phospholipid synthesis could also compensate for LPS depletion^16,27,36^. Reduced fatty acid synthesis could slow expansion of the inner membrane and thereby better match expansion of the LPS-depleted OM, whereas increased phospholipid synthesis could potentially increase phospholipid delivery to the OM. We therefore altered activity of *fadR*, a major regulator of fatty acid metabolism. Both deletion and overexpression of *fadR* from an IPTG-inducible promoter (*fadR*::*kan* and p*fadR*, respectively) reduced the CHIR-090 ZOI (Δ*fadR*, 11.9% smaller than wild type; p*fadR*, 5.7% smaller than the empty-vector control; Table S4). Thus, perturbing phospholipid metabolism in either direction can improve growth in the presence of CHIR-090, indicating that susceptibility to LPS depletion depends on phospholipid homeostasis rather than simply on the absolute rate of phospholipid synthesis.

Together, these results show that perturbations affecting phospholipid synthesis and distribution can partially compensate for LPS depletion. Combined with the failure of osmoprotection alone to preserve culturability, these findings support a role for LPS in coordinating membrane biogenesis that is distinct from its contribution to cell-envelope mechanics.

### Mutations predicted to increase cell-envelope rigidity partially compensate for LPS depletion

Although osmoprotection did not preserve culturability, its strong protection of cell-envelope integrity suggested that increasing cell wall rigidity might still partially compensate for LPS depletion. We therefore predicted that mutations that increase peptidoglycan crosslinking should improve growth during LPS depletion. Consistent with this possibility, a mutation affecting the transpeptidase PBP1a that increases 3,3-peptidoglycan crosslinks promotes growth of LPS-deficient *A. baumannii*^23,24^. To test this prediction in *E. coli*, we screened mutants predicted to alter peptidoglycan remodeling using the CHIR-090 Kirby-Bauer assay.

Mutations predicted to increase peptidoglycan crosslinking reduced susceptibility to CHIR-090. Of the thirteen peptidoglycan-related mutants examined, deletion of the D,D-endopeptidase *dacB* (*dacB*::*kan*) and deletion of all three amidases (Δ*amiABC*) (*amiA*::*frt amiB*::*kan amiC*::*frt*) reduced CHIR-090 ZOIs by 3.9% and 11.6%, respectively, relative to the parental strain (Table S4, Fig 3B). Because DacB and the amidases cleave peptidoglycan during cell wall remodeling, loss of these activities is predicted to increase peptidoglycan crosslinking and mechanical rigidity. CHIR-090 depleted LPS to a similar extent in *ΔamiABC* and parental cells (Fig 3D), indicating that its reduced susceptibility was not due to preservation of LPS abundance. As observed for Δ*pldA*, high osmolarity slowed the loss of culturability of Δ*amiABC* following CHIR-090 treatment (Fig 3D), suggesting that increased cell wall rigidity and osmoprotection provide complementary forms of mechanical protection during LPS depletion.

In contrast to observations in *A. baumannii*^15,23^, deletion of the *E. coli* PBP1a homolog (*mrcA*::*kan*) had no detectable effect on CHIR-090 susceptibility (Table S4), indicating that the specific peptidoglycan perturbations that compensate for LPS depletion differ between species. Together, these findings provide additional evidence that LPS contributes to the mechanical integrity of the cell envelope and demonstrate that increasing peptidoglycan rigidity can partially compensate for its depletion.

### A gain-of-function mutation in the cell division protein FtsA improves growth during LPS depletion

LPS depletion disrupts cell division, although the underlying mechanism remains unclear^11,12,37^. We therefore reasoned that if impaired divisome activity contributes directly to growth arrest during LPS depletion, perturbations that enhance division should improve growth. To test this prediction, we examined gain-of-function alleles and overexpression constructs known to promote divisome assembly or activation, including *ftsA*(R286W) (hereafter *ftsA\**), *ftsL*(E88K), *ftsW*(M269I), *ftsW*(E289G), and overexpression of *ftsN* (p*ftsN*, under an L-arabinose-inducible promoter) or *ftsQAZ* (p*ftsQAZ*, under the native promoter). Each of these perturbations enhances divisome assembly or activation and reduces cell size^38–40^.

Of the six perturbations examined, only *ftsA\** improved growth in the presence of CHIR-090, reducing the ZOI by 15.3% (Fig 3B, Table S4). In contrast, overexpression of *ftsQAZ* slightly increased CHIR-090 susceptibility, while the remaining perturbations had no detectable effect (Table S4). Thus, enhancing divisome activity does not generally compensate for LPS depletion. CHIR-090 depleted LPS to a similar extent in *ftsA\** and parental cells (Fig 3E), indicating that its reduced susceptibility was not due to preservation of LPS during treatment. Moreover, *ftsA\** did not preserve culturability during prolonged LPS depletion, with CFUs declining similarly to those of the parental strain under both standard- and high-osmolarity conditions (Fig 3E).

Notably, *ftsA*\* cells contained ∼1.7-fold more LPS than wild-type cells before CHIR-090 treatment (Dunnett’s test*, p* = 0.025; Fig 3E and 1B). Together, these results argue against impaired divisome activity as a general explanation for growth arrest during LPS depletion. Instead, the unique phenotype of *ftsA\**, together with its elevated basal LPS abundance, raises the possibility of a more specific connection between FtsA and LPS homeostasis.

### Spontaneous mutations strongly protect against LPS depletion

To identify mechanisms that compensate for LPS depletion beyond those captured by our candidate-gene approach, we isolated spontaneous CHIR-090-resistant colonies arising within ZOIs in *E. coli* and *K. pneumoniae* (Fig 3A). Among five *K. pneumoniae* isolates, four carried *lpxC*(V37G), an allele previously associated with resistance to LpxC inhibition^27^. This allele increases cellular LpxC abundance, likely explaining the resistance phenotype. The fifth isolate carried a frameshift in *fadR*, consistent with our finding that *fadR* deletion reduces CHIR-090 susceptibility in *E. coli* (Fig 3B).

Of six *E. coli* isolates, four exhibited heritable CHIR-090 resistance (Fig 4A). Two carried an identical in-frame 6-bp deletion in *marR*, removing amino acids 31 and 32 (hereafter *marR*^Δ^^31–32^). The remaining two carried an *IS421* insertion in the *lon* promoter (*P_lon_*::*IS421*) together with a nonsense mutation in *plsX* [*plsX*(L25*)]. *marR* encodes the transcriptional repressor of the *<u>m</u>*ultiple *<u>a</u>*ntibiotic *<u>r</u>*esistance (*marRAB)* operon^41^, whereas Lon is a major ATP-dependent protease involved in protein homeostasis^42^ and PlsX participates in phospholipid metabolism^43^. None of these mutations had previously been implicated in CHIR-090 resistance in *E. coli*, prompting us to determine how they protect against LPS depletion.

**Figure 4:**
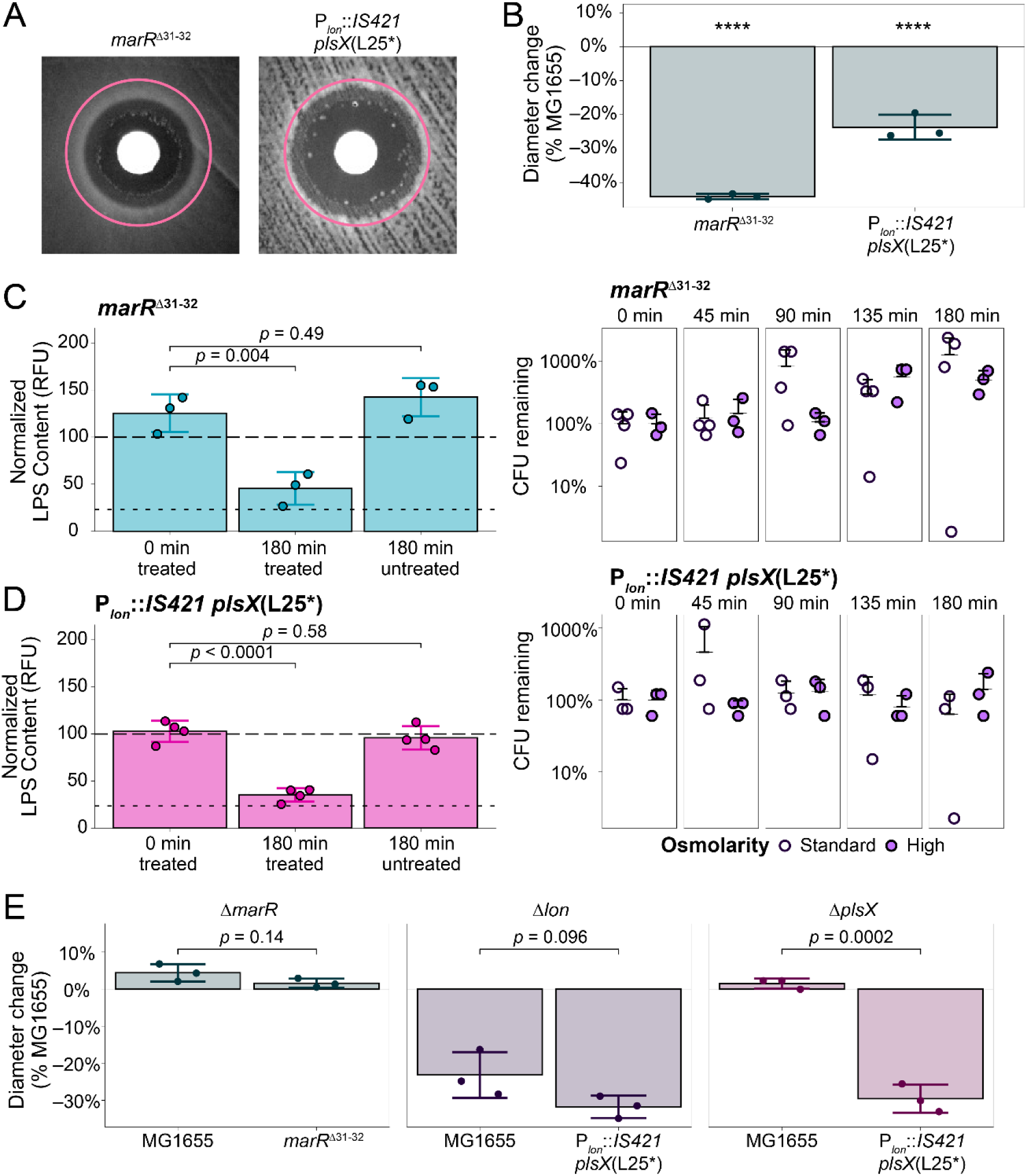
Spontaneous mutations in *marR* and *lon* preserve culturability during LPS depletion. **(A)** Representative ZOIs produced by CHIR-090 in the spontaneous *marR*^Δ^^31–32^ and *P_lon_*::*IS421 plsX*(L25*) isolates. Pink circles indicate the ZOI diameter of wild-type MG1655 (27 mm). **(B)** Percent change in ZOI diameter relative to MG1655 for the two spontaneous isolates, quantified as in Fig 3B. Both isolates exhibited substantially reduced ZOIs, indicating increased growth in the presence of CHIR-090. ****, *p*<0.0001. **(C,D)** Relative LPS abundance (left) and culturability (right) during CHIR-090 treatment of the *marR*^Δ^^31–32^ (C) and *P_lon_*::*IS421 plsX*(L25*) (D) isolates. Both isolates maintained culturability despite substantial LPS depletion. LPS abundance and culturability were quantified as in Fig 3C-E and 2E-G, respectively. Dashed and dotted lines indicate the mean pretreatment and 180-min CHIR-090-treated LPS abundance, respectively, for MG1655. **(E)** Effects of introducing Δ*marR*, Δ*lon*, or Δ*plsX* into the indicated backgrounds on CHIR-090 ZOI diameter. Values are shown relative to MG1655. The *marR*^Δ^^31–32^ phenotype was not reproduced by Δ*marR*, whereas Δ*lon* phenocopied the reduced CHIR-090 susceptibility of the *P_lon_*::*IS421 plsX*(L25*) isolate and Δ*plsX* did not, implicating the *lon* promoter insertion rather than *plsX*(L25*) as the causal mutation. Each point represents a biological replicate. Bars and error bars indicate mean ± SD. *p*-values were determined using Dunnett’s multiple-comparisons test.

Among all mutants examined, *marR*^Δ^^31–32^ conferred the strongest protection against CHIR-090, reducing the ZOI by 44% relative to the parental strain (Fig 4A, B). The deleted residues lie near the MarR DNA-binding domain^44^, suggesting that the mutation could alter transcriptional regulation. Deletion of *marR* (*marR*::*kan*) did not reproduce the *marR*^Δ^^31–32^ phenotype, whereas expression of wild-type *marR* in trans restored CHIR-090 susceptibility of *marR*^Δ^^31–32^ cells to that of the parental strain (Fig 4E, Table S4). These results indicate that *marR*^Δ^^31–32^ is not simply a loss-of-function allele and that its phenotype is suppressed by wild-type MarR.

For the second spontaneous *E. coli* genotype, *P_lon_*::*IS421* rather than *plsX*(L25*) was responsible for reduced CHIR-090 susceptibility. Deletion of *lon* largely phenocopied the spontaneous isolate (Fig 4E), whereas deletion of *plsX* (*plsX*::*kan*) had no detectable effect on CHIR-090 susceptibility either alone or in the *P_lon_*::*IS421* background (Fig 4E). We therefore refer to this isolate hereafter by the causative mutation, *P_lon_*::*IS421*.

Unlike the candidate-gene perturbations examined above, both *marR*^Δ^^31–32^ and *P_lon_*::*IS421* strongly preserved culturability during prolonged LPS depletion. CHIR-090 depleted LPS to a similar extent in both mutants and parental cells (Fig 4C,D), indicating that their protection was not explained by preservation of LPS abundance. Nevertheless, CFUs of *marR*^Δ^^31–32^ increased during CHIR-090 treatment, while those of *P_lon_*::*IS421* remained approximately constant, in striking contrast to the rapid decline in culturability of parental cells (Fig 4C, D). Thus, spontaneous mutations in *marR* and *lon* enable *E. coli* to remain culturable despite substantial LPS depletion, revealing compensatory mechanisms distinct from maintaining LPS abundance.

### Increased MarA activity promotes growth during LPS depletion

MarR and Lon negatively regulate MarA through distinct mechanisms: MarR represses *marA* transcription, whereas the ATP-dependent protease Lon degrades MarA^42,45,46^. Thus, the independent isolation of mutations affecting *marR* and *lon* suggested that increased MarA activity might underlie their protection against LPS depletion. Consistent with this model, *marR*^Δ^^31–32^ increased *marA* transcript abundance ∼69-fold (Table 1). In contrast, *marA* transcript abundance increased only ∼1.5-fold in both *P_lon_*::*IS421* and Δ*lon* cells (Table 1), as expected if loss of Lon increases MarA activity primarily through post-transcriptional stabilization.

**Table 1:** Relative transcript abundance of *marA*, *acrB*, and *tolC* measured by RT-qPCR.

| Strain/condition | *marA* fold-change | *acrB* fold-change | *tolC* fold-change |
| --- | --- | --- | --- |
| MG1655 | 1.0 ± 0.1 | 1.0 ± 0.04 | 1.0 ± 0.2 |
| MG1655 (CHIR-090 treated) | 2.2 ± 0.2 | 1.8 ± 0.3 | 0.9 ± 0.1 |
| pEV | 0.8 ± 0.1 | 1.0 ± 0.01 | 1.0 ± 0.1 |
| *pmarA* | 358 ± 48 | 1.6 ± 0.2 | 2.3 ± 0.1 |
| *marR*<sup>Δ31-32</sup> | 69 ± 12 | 2.2 ± 0.1 | 3.4 ± 0.4 |
| *P<sub>lon</sub>::IS421* | 1.5 ± 0.2 | 1.5 ± 0.3 | 1.6 ± 0.3 |
| *Δlon* | 1.5 ± 0.2 | 1.5 ± 0.2 | 1.6 ± 0.4 |

Values represent mean fold-change ± SD from at least three biological replicates.
Transcript abundance was normalized to the reference genes *gyrA* and *rpsL* and is shown relative to untreated MG1655.

To determine whether increased MarA activity is sufficient to promote growth during tolerance to LPS depletion, we expressed *marA* from an IPTG-inducible promoter (p*marA*). Induction increased *marA* transcript abundance ∼360-fold relative to wild-type cells (Table 1). MarA overexpression reduced the CHIR-090 ZOI by ∼30% relative to the empty-vector control (Fig 5A). CHIR-090 nevertheless depleted LPS to ∼25% of its initial abundance in MarA-overexpressing cells (Fig 5B), indicating that protection was not explained by preservation of LPS abundance. MarA overexpression also improved growth during *lpxC* tCRISPRi-mediated depletion (Fig 5C), demonstrating that its protective effect was independent of the method used to inhibit LPS synthesis.

**Figure 5:**
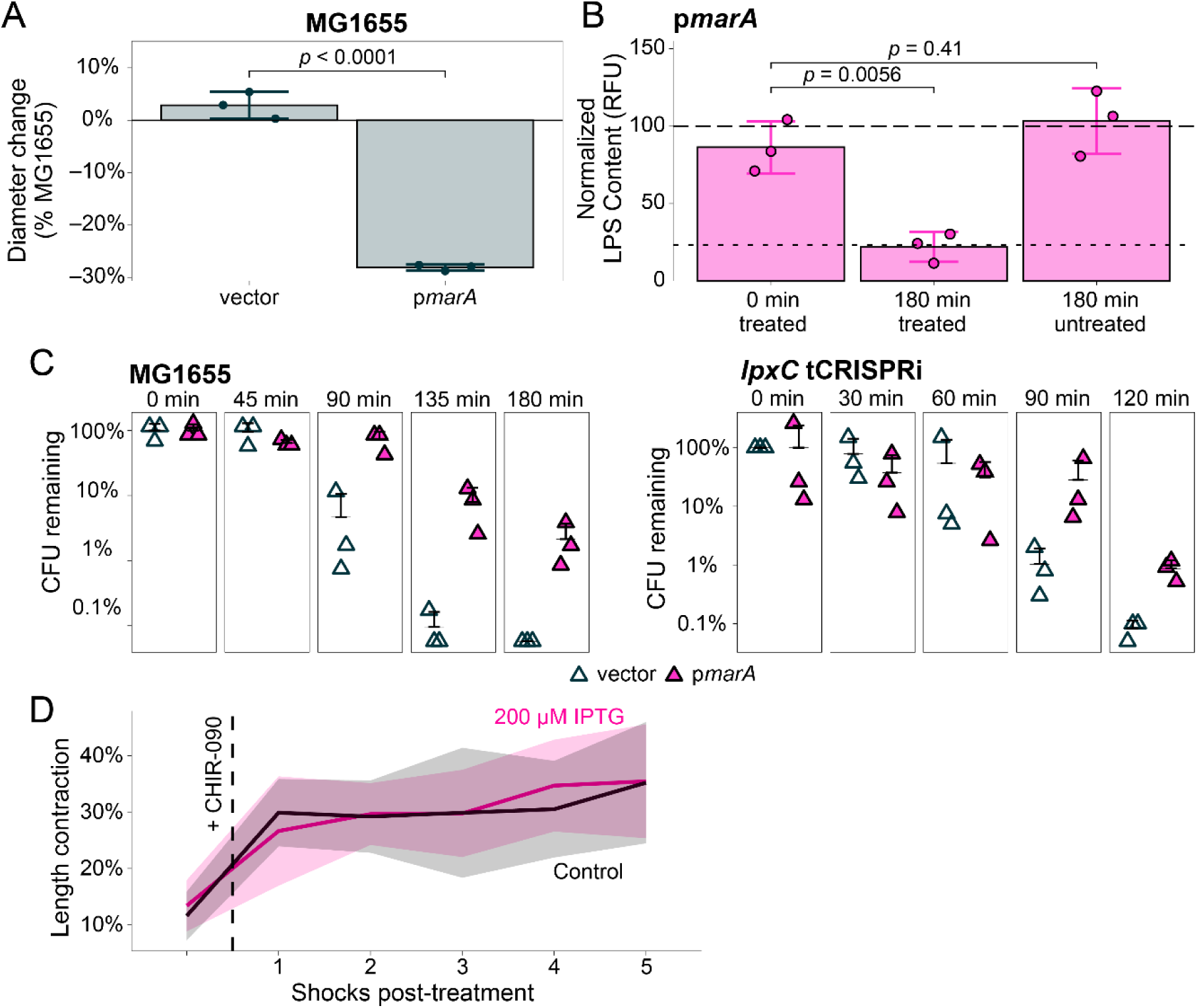
MarA promotes growth during LPS depletion without restoring LPS abundance or cell-envelope stiffness. **(A)** Percent change in CHIR-090 ZOI diameter for *E. coli* carrying p*marA* or the empty-vector control, relative to MG1655. *marA* overexpression substantially reduced the ZOI, indicating increased growth in the presence of CHIR-090. **(B)** Relative LPS abundance before and after 180 min of CHIR-090 treatment in cells carrying p*marA*, quantified as in Fig 3C-E. LPS abundance decreased despite *marA* overexpression, indicating that MarA does not protect by preventing LPS depletion. Dashed and dotted lines indicate the mean pretreatment and 180-min CHIR-090-treated LPS abundance, respectively, for MG1655. **(C)** Culturability during LPS depletion in MG1655 and *lpxC* tCRISPRi cells carrying p*marA* or the empty-vector control. LPS depletion was induced with CHIR-090 in MG1655 or by L-arabinose induction in the *lpxC* tCRISPRi strain. *marA* overexpression preserved culturability with both methods of LPS depletion. CFU/mL at each time point is shown relative to the corresponding *t*_0_ value. **(D)** Fractional cell-length contraction during successive hyperosmotic shocks in cells carrying p*marA* with or without IPTG induction. *marA* overexpression did not detectably alter the mechanical response to LPS depletion, indicating that its protective effect does not require restoration of cell-envelope stiffness. CHIR-090 was added after the pretreatment shock, as indicated by the dashed line. Lines and shaded regions indicate mean ± SD. Each point in (A-C) represents a biological replicate; bars and error bars indicate mean ± SD. *p*-values were determined using Dunnett’s multiple-comparisons test.

We next asked whether MarA protects cells by compensating for the loss of OM mechanical rigidity. Reduced LPS abundance decreases OM stiffness, resulting in greater cell shrinkage following hyperosmotic shock^18^. If MarA acts by restoring OM rigidity, its overexpression should reduce this response during LPS depletion.

MarA overexpression did not detectably restore OM rigidity. CHIR-090 treatment increased cell shrinkage during repeated hyperosmotic shocks (Fig 5D), consistent with a rapid reduction in OM stiffness. However, CHIR-090-treated p*marA* cells exhibited similar shrinkage in the presence and absence of IPTG (Fig 5D). Thus, increased MarA activity promotes growth during LPS depletion without restoring LPS abundance or OM mechanical rigidity, pointing to a protective mechanism distinct from the mechanical function of LPS.

### AcrAB-TolC is required for MarA*-*mediated protection during LPS depletion

The genes encoding the AcrAB-TolC multidrug efflux pump are among the best-characterized members of the MarA regulon^41,47^. AcrAB-TolC spans the cell envelope^48^ and confers resistance to a broad range of structurally diverse antibiotics^49^. RT-qPCR analysis indicated that *acrB*, used as a proxy for the *acrAB* operon, and *tolC* were modestly upregulated in each of the MarA-activating backgrounds (Table 1). *acrB* transcript abundance increased 2.2-fold in *marR*^Δ^^31–32^, 1.5-fold in *P_lon_*::*IS421*, and 1.4-fold following *marA* overexpression, while *tolC* increased 3.4-fold, 1.5-fold, and 2.3-fold, respectively.

We therefore asked whether AcrAB-TolC is required for MarA-mediated protection during LPS depletion. Deletion of *acrB* (*acrB*::*kan*), or *tolC* (*tolC*::*kan*) abolished the reduction in CHIR-090 susceptibility conferred by *marA* overexpression, while deletion of *acrA* substantially reduced it (Fig 6A). Similarly, deletion of *acrA* or *acrB* eliminated the reduced CHIR-090 susceptibility of both *marR*^Δ^^31–32^ and *P_lon_*::*IS421* spontaneous mutants (Fig 6B). Thus, AcrAB-TolC is required for protection conferred by multiple independent perturbations that increase MarA activity.

**Figure 6:**
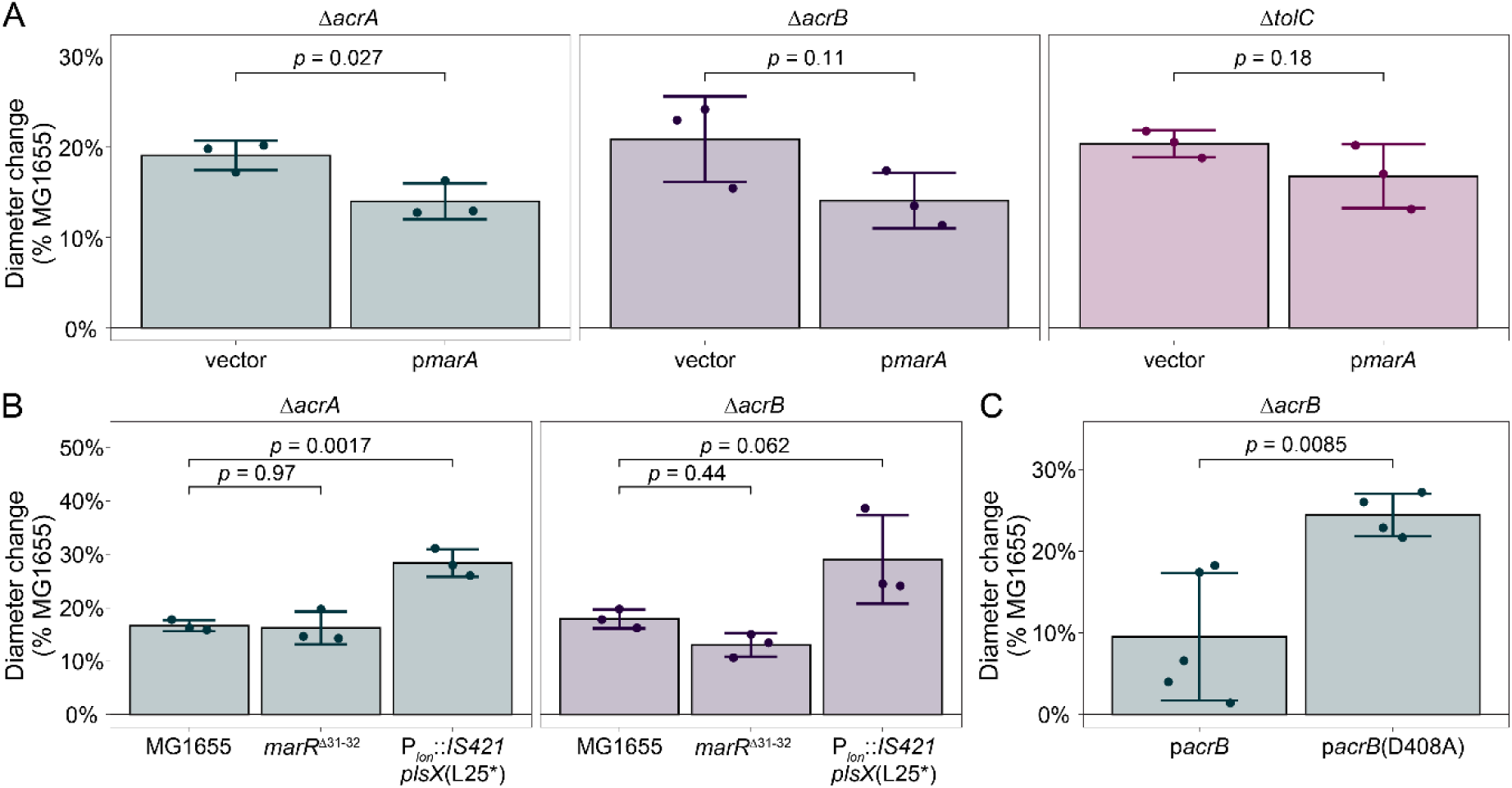
AcrAB-TolC is required for MarA-mediated protection during LPS depletion. **(A)** Percent change in CHIR-090 ZOI diameter in Δ*acrA*, Δ*acrB*, and Δ*tolC* strains carrying p*marA* or the empty-vector control, relative to MG1655. Deletion of each AcrAB-TolC component reduced or eliminated the protective effect of *marA* overexpression. **(B)** Percent change in CHIR-090 ZOI diameter following deletion of *acrA* or *acrB* in MG1655, *marR*^Δ^^31–32^, and *P_lon_*::*IS421 plsX*(L25*) backgrounds. Loss of *acrA* or *acrB* also reduced the protection conferred by the spontaneous suppressors, consistent with their effects being mediated through AcrAB-TolC. **(C)** Percent change in CHIR-090 ZOI diameter in Δ*acrB* cells expressing wild-type AcrB or the transport-defective AcrB(D408A) variant. AcrB(D408A) failed to restore the protection provided by wild-type AcrB, demonstrating that AcrB transport activity, rather than simply its presence in the envelope, is required for protection during LPS depletion. Each point represents a biological replicate; bars and error bars indicate mean ± SD. *p*-values were determined using Dunnett’s multiple-comparisons test.

CHIR-090 treatment itself also activated components of the MarA response in wild-type cells, suggesting that this pathway responds to LPS depletion even in the absence of genetic perturbation. After 90 min of CHIR-090 treatment, *marA* and *acrB* transcript abundance increased 2.2-fold and 1.8-fold, respectively, whereas *tolC* expression was unchanged (Table 1). Thus, LPS depletion induces *marA* and *acrAB* expression, while the spontaneous suppressor mutations amplify a pathway whose activity contributes to growth during LPS depletion.

### AcrAB-TolC transport activity promotes growth during LPS depletion

We next asked whether the requirement for AcrAB-TolC during LPS depletion reflects its transport activity or a transport-independent structural or regulatory role within the cell envelope. To distinguish between these possibilities, we used the previously characterized efflux-deficient *acrB*(D408A) allele^50^. AcrB is the inner membrane component of AcrAB-TolC and captures substrates primarily from the periplasm and inner membrane-periplasm interface^51^. Transport is driven by the proton motive force through a proton relay within AcrB^52^. The D408A substitution disrupts a critical proton-binding residue required for transport without preventing assembly of the efflux complex^53^.

We expressed either wild-type *acrB* or *acrB*(D408A) from IPTG-inducible plasmids [p*acrB* and p*acrB*(D408A), respectively] in Δ*acrB* cells. Deletion of *acrB* increased the CHIR-090 ZOI by 16.3% relative to wild-type cells (Table S4). Expression of wild-type *acrB* restored CHIR-090 susceptibility to wild-type levels, whereas *acrB*(D408A) failed to complement Δ*acrB* (Fig 6C). Thus, the transport activity of AcrAB-TolC, rather than simply the presence of the efflux complex, is required to promote growth during LPS depletion.

## Discussion

LPS is integrated into numerous aspects of Gram-negative cell envelope physiology, making it difficult to identify a single property that explains why it is essential in most Gram-negative bacteria. Our findings suggest that this difficulty reflects the biology itself: LPS essentiality does not appear to arise from one indispensable activity. Instead, LPS contributes to multiple features of the envelope that become difficult to maintain simultaneously when LPS synthesis is restricted. This view helps reconcile the diverse phenotypes associated with LPS depletion and the observation that perturbations affecting seemingly distinct processes can each provide partial protection without eliminating the requirement for LPS.

One important contribution is mechanical^18^. The ability of osmoprotection to largely preserve envelope integrity without preserving culturability demonstrates that catastrophic envelope failure is not the sole reason that LPS-depleted cells cease growing. Conversely, the protective effects of perturbations predicted to increase peptidoglycan rigidity support the importance of LPS to envelope mechanics. LPS therefore appears to occupy an unusual position in the envelope: its mechanical properties are important, but preserving those properties alone is insufficient to sustain growth. This distinguishes LPS depletion from perturbations such as loss of the cell wall, for which appropriate osmotic conditions can permit continued proliferation as L-forms.

A second challenge is likely created by the physical requirement to build an OM as cells grow. LPS constitutes a major fraction of the outer leaflet, so restricting its availability creates a problem of membrane occupancy even if membrane rupture can be prevented. The effects of perturbing *pldA*, *mlaA*, and *fadR* are consistent with the idea that altering phospholipid homeostasis can partially accommodate this imbalance. Similar genetic adaptations in LPS-deficient *A. baumannii* suggest that maintaining appropriate membrane composition may represent a general challenge created by LPS depletion. Nevertheless, phospholipids are unlikely to be complete substitutes for LPS, which also influences OM protein assembly^54,55^, membrane organization, and fluidity^19^. Thus, the requirement for LPS may arise partly because membrane composition, mechanics, and protein organization cannot be varied independently.

This framework may also explain why the genetic routes that compensate for LPS depletion differ between organisms. Increased peptidoglycan crosslinking is associated with protection in both *E. coli* and *A. baumannii*^24^, yet the specific perturbations that produce this effect are not interchangeable between species. The underlying physiological constraints imposed by LPS depletion may therefore be conserved even when the architecture of the compensatory response is not. Such differences could help explain the striking ability of a small number of Gram-negative species, including *A. baumannii*, to evolve complete LPS independence while LPS remains essentially indispensable in most others.

By contrast, our results provide little support for defective cell division as a general explanation for LPS essentiality. The exceptional behavior of *ftsA\** is more suggestive of a specific connection between FtsA and envelope homeostasis than of rescue through enhanced cytokinesis, particularly given the elevated basal LPS abundance in this mutant. Determining whether this reflects direct coordination between divisome activity and LPS biogenesis will require further study, but the result emphasizes that cellular processes associated with LPS depletion need not themselves explain why LPS is essential.

The identification of the MarA–AcrAB-TolC pathway provides a different perspective on how cells accommodate LPS depletion. Rather than preserving LPS abundance or restoring the mechanical properties of the OM, increased MarA activity allows cells to better tolerate the physiological consequences of having less LPS. The convergence of independent *marR* and *lon* suppressors on MarA, together with the induction of *marA* and *acrAB* by CHIR-090 itself, suggests that the suppressors amplify an endogenous envelope-stress response rather than creating an entirely new cellular state. Most importantly, protection requires ArcAB-TolC transport activity, implicating movement of one or more substrates across the envelope in adaptation to LPS depletion.

The identity of the relevant substrate is therefore an important question. One attractive possibility is that AcrAB-TolC contributes directly to lipid homeostasis. Reconstituted AcrAB complexes can transport phospholipids^56^, and AcrAB-TolC has been implicated genetically in free-fatty-acid export^57^. Export of excess phospholipids or related lipid species could provide a means of accommodating continued inner-membrane lipid synthesis when LPS availability constrains expansion of the outer leaflet. This model would place AcrAB-TolC within the same broader physiological framework suggested by the *pldA*, *mlaA*, and *fadR* phenotypes: survival during LPS depletion may depend in part on the ability to rebalance lipid flux across the envelope. A connection between efflux and phospholipid homeostasis has also been reported in *A. baumannii*^58^, although direct measurement of the substrates transported by AcrAB-TolC during LPS depletion will be necessary to test this model.

More generally, this possibility suggests that multidrug efflux pumps may have important physiological functions that are obscured by their prominent role in antimicrobial resistance. AcrAB-TolC is generally considered a defense against exogenous toxic compounds^59^, but its broad substrate specificity and position spanning the cell envelope make it well suited to participate in envelope remodeling. The observation that increased expression of the related MexAB-OprM pump protects *Pseudomonas aeruginosa* from CHIR-090^60^ raises the possibility that this function extends beyond *E. coli*. Endogenous substrate transport could therefore represent an underappreciated role of multidrug efflux systems, particularly when membrane composition or biogenesis is perturbed.

Together, our findings suggest that LPS essentiality is best understood as an emergent property of its integration into the Gram-negative cell envelope rather than as the consequence of a single indispensable biochemical activity. This framework explains why compensating for individual consequences of LPS depletion can improve growth without eliminating the requirement for LPS. It also raises an evolutionary possibility: LPS need not have originated as an essential molecule. Once incorporated into the OM, its physical and chemical properties could have been progressively exploited by other envelope processes, making those processes increasingly dependent on its presence. In this view, essentiality can evolve not because a molecule acquires one uniquely indispensable activity, but simply because cellular physiology becomes organized around it.

## Methods

### Strains, plasmids, and growth conditions

Bacterial strains, plasmids, and oligonucleotides used in this study are listed in Tables S1, S2, and S3, respectively. Unless otherwise specified, strains were cultivated in EZ Rich Defined Medium (Teknova) at 37 °C with shaking at 200 rpm. For plasmid maintenance, antibiotics were added at the concentrations indicated in Table S2. Because glucose or other fermentable carbon sources can interfere with arabinose-dependent regulation of the *lpxC* sgRNA cassette, tCRISPRi strains were grown in medium supplemented with 0.2% (v/v) glycerol as the sole carbon source. High-osmolarity conditions were achieved by supplementing the indicated growth medium with 500 mM D-sorbitol. tCRISPRi strains were generated by electroporation using the oligonucleotides listed in Table S3 according to a previously established protocol^61^.

### MIC assays

Minimum inhibitory concentrations (MICs) were determined by broth microdilution. Bacterial cultures were grown to mid-exponential phase (OD_600_∼0.2–0.6) in LB medium and diluted to ∼10^5^ CFU/mL into fresh medium containing two-fold serial dilutions of the indicated antimicrobial agent in sterile flat-bottom 96-well plates and incubated at 37 °C with shaking at 200 rpm for 20 h. The MIC was defined as the lowest antimicrobial concentration that completely prevented visible growth relative to the untreated control.

### Growth kinetics

Cultures were grown from single colonies to mid-exponential phase (OD_600_∼0.2–0.6) under standard conditions, pelleted, and resuspended in prewarmed medium to OD_600_=1 (∼10^9^ cells/mL). Propidium iodide (PI; 1.5 µM) was included to match conditions used for time-lapse microscopy. For inhibitor treatments, CHIR-090 was added at 0.25 µg/mL for *E. coli* MG1655 or 1 µg/mL for *K. pneumoniae* TOP52, corresponding to ∼3X the empirically determined MIC for each strain. For tCRISPRi experiments, L-arabinose was added to 0.2% (w/v) to induce *lpxC* sgRNA expression.

Cell suspensions were serially diluted 1:10 and 1:100 in medium containing the corresponding treatment, and 150-µL aliquots of the undiluted resuspension and each dilution were transferred into black, clear-bottom 96-well plates. Plates were sealed with gas-permeable membranes and incubated at 37 °C for 20 h on a Synergy H1 microplate reader (BioTek) with orbital shaking at 232 cpm. OD_600_ was measured every 10 min.

### Lipopolysaccharide quantification

Cultures were grown to mid-exponential phase (OD_600_∼0.2–0.6) under standard conditions. An aliquot was harvested at *t*_0_ and stored at –80 °C for baseline comparison. Remaining cells were diluted to OD_600_=0.01 in fresh medium supplemented with either ∼3X the empirically determined wild-type MIC of CHIR-090 (0.25 µg/mL for *E. coli* MG1655 or 1 µg/mL for *K. pneumoniae* TOP52) or 0.2% (w/v) L-arabinose for tCRISPRi induction. Cultures were incubated at 37 °C for 180 min, after which cells were harvested by centrifugation and stored at –80 °C.

Cell pellets were thawed, resuspended in 50 µL of B-PER Bacterial Protein Extraction Reagent (Thermo Fisher Scientific), and incubated at room temperature for 15 min. Insoluble material was removed by centrifugation at 2150*g* for 5 min at room temperature. Total protein concentration in the clarified lysates was determined using a Pierce BCA Protein Assay Kit (Thermo Fisher Scientific) according to the manufacturer’s microplate protocol. Lysates were digested with Proteinase K (1 µL of 20 mg/mL Proteinase K per 30 µL sample; Thermo Fisher Scientific) at 55 °C for ∼16 h to remove contaminating proteins.

Samples were normalized by total protein content, mixed with modified 4X Laemmli buffer (40% (w/v) sucrose, 4% (w/v) sodium dodecyl sulfate, 1.97% (w/v) Tris-HCl, 0.015% (w/v) bromophenol blue in ultrapure H_2_O) to a final concentration of 1X, and supplemented with 2-mercaptoethanol to a final concentration of 10% (v/v). Glycerol in standard Laemmli buffer was replaced with sucrose to minimize potential interference with downstream staining. Samples were heated at 95 °C for 5 min, and equal protein loads were separated by SDS-PAGE on precast 4–20% polyacrylamide gradient gels (Bio-Rad) using Tris-glycine-SDS running buffer. Gels were stained for 120 min using the Pro-Q Emerald 300 Lipopolysaccharide Gel Staining Kit (Thermo Fisher Scientific) according to the manufacturer’s instructions and imaged using a Bio-Rad Gel Doc EZ Imager. LPS band intensities were quantified using FIJI^44^.

To account for variation in staining between gels, LPS band intensities were normalized to the 42-kDa band of the Candycane Glycoprotein Molecular Weight Standards (Thermo Fisher Scientific) included on each gel. Values were additionally normalized to the amount of total protein loaded and scaled such that 100 represented the mean *t*_0_ value for *E. coli* MG1655 grown under standard-osmolarity conditions. Total protein loaded was kept consistent across time points of each experiment but varied slightly for different strains and medium conditions to accommodate differences in growth rate during the assay.

### Time-lapse microscopy

Cultures were grown to early exponential phase (OD_600_∼0.1–0.2) under standard conditions, harvested, and mounted on 1% agarose pads prepared with the corresponding growth medium containing 1.5 µM PI and either ∼3X the empirically determined wild-type MIC of CHIR-090 (0.25 µg/mL for *E. coli* MG1655 or 1 µg/mL for *K. pneumoniae* TOP52) or 0.2% (w/v) L-arabinose for tCRISPRi induction. Samples were allowed to equilibrate on the pads for ∼5 min until excess liquid was absorbed; drying time was minimized to limit changes in local osmolarity.

Imaging was performed using a Nikon Ti-E or Ti2-E inverted microscope (Nikon Instruments). The Ti-E was equipped with a 100X Plan N Ph3 objective (NA 1.45), an X-Cite 120 LED illuminator (Lumen Dynamics), and an ORCA-Flash 4.0 V2 camera (Hamamatsu Photonics). The Ti2-E was equipped with an identical objective, a Lumencor SOLA LED light engine (Lumencore), and an ORCA-Fusion sCMOS camera (Hamamatsu Photonics). PI fluorescence was imaged using Chroma AT560/40x and AT635/60m excitation and emission filters, respectively, and an AT600DC dichroic mirror (Chroma Technology Corporation). The microscope stage was enclosed in an environmental chamber (Okolab) maintained at 37 °C. Images were acquired using NIS-Elements Advanced Research software.

Cell envelope integrity was assessed by PI staining. Cells exhibiting uniform PI staining were classified as having lost membrane barrier integrity, and the proportion of PI-positive cells was quantified using FIJI^62^.

### Culturability assays

To assess culturability during LPS depletion, cultures were grown in LB medium to mid-exponential phase (OD_600_∼0.2–0.6), diluted to OD_600_=0.01 in fresh medium containing ∼3X the empirically determined wild-type MIC of CHIR-090 (0.25 µg/mL for *E. coli* MG1655 or 1 µg/mL for *K. pneumoniae* TOP52), and incubated at 37 °C with shaking. For tCRISPRi strains, cells were grown in EZ Rich Defined Medium containing 0.2% (v/v) glycerol and induced with 0.2% (w/v) L-arabinose.

Aliquots were collected every 45 min for up to 180 min during CHIR-090 treatment and every 30 min for up to 120 min following tCRISPRi induction. Samples were serially diluted 10-fold (10^−1-10–5^) in the corresponding growth medium, and a defined volume of each dilution was plated on LB agar. Plates were incubated at 37 °C for ∼16 h before colonies were counted. Culturability was expressed as CFU/mL at each time point relative to the corresponding *t*_0_ value. Where indicated, 500 mM D-sorbitol was included in both the culture medium and LB agar to maintain consistent osmolarity.

### Candidate gene screen

To identify mutations that alter susceptibility to CHIR-090, we used a modified Kirby-Bauer disk diffusion assay^35^. Bacterial cultures were grown to OD_600_∼0.1, and ∼200 µL of culture were uniformly spread onto prewarmed LB agar plates (90-mm diameter; 25 mL of LB agar per plate) to generate a dense lawn. An 8-mm Whatman Grade 1 filter-paper disk containing 1 µg of CHIR-090 in dimethyl sulfoxide was allowed to dry and then placed at the center of each plate. Plates were incubated at 37 °C for 20 h and imaged using a Bio-Rad Gel Doc EZ Imager. ZOI diameters were measured using FIJI^62^.

### Genomic analysis of spontaneously resistant isolates

To identify mutations associated with reduced CHIR-090 susceptibility, colonies emerging at the inner periphery of the ZOI were isolated from wild-type *E. coli* MG1655 and *K. pneumoniae TOP52* lawns generated during the candidate-gene screen and submitted to SeqCenter for processing. Genomic DNA was extracted using a ZymoBIOMICS DNA Miniprep Kit according to the manufacturer’s protocol, with ∼50–100 mg of cell pellet resuspended in 750 µL of lysis buffer. Sequencing libraries were prepared using the Illumina DNA Prep Kit and multiplexed using custom 10-bp dual indexes (Integrated DNA Technologies). Library fragments were targeted to ∼280 bp without additional fragmentation or size selection. Libraries were sequenced on an Illumina NovaSeq X Plus platform using 2×151 bp paired-end reads.

Sequencing reads were demultiplexed, quality filtered, and adapter trimmed using bcl-convert v4.2.2 (Illumina). Reads were aligned to the corresponding reference genome (*E. coli* MG1655, NCBI accession GCF_904425475; *K. pneumoniae* TOP52, NCBI accession GCF_003957455.1_ASM395745v1), and variants were identified using breseq v0.39.0 with default parameters^63^.

### Cell-envelope stiffness measurements

Cell-envelope stiffness was assessed using a microfluidic osmotic shock assay based on a previously described plasmolysis assay^18^. Exponentially growing cells were loaded into a CellASIC ONIX microfluidic flow cell (Sigma-Aldrich, Cat. #B04A-03-5PK) and maintained at 37 °C using an environmental chamber (HaisonTech). Medium exchanges were controlled using a CellASIC ONIX2 microfluidic platform (Sigma-Aldrich, Cat. #CAX2-S0000). Cells were subjected to repeated hyperosmotic shocks at 35-min intervals, consisting of 5 min in LB supplemented with 1 M sorbitol (Sigma-Aldrich, Cat. #S6021), followed by 30 min in LB at standard osmolarity. Twenty cells were analyzed per condition per osmotic-shock cycle.

Phase-contrast and fluorescence images were acquired throughout the assay using a Nikon Eclipse Ti-E inverted microscope equipped with a 100× oil-immersion objective (NA 1.40; Nikon Instruments) and a Prime BSI Express sCMOS camera (Teledyne Photometrics). Cell contours were segmented using DeepCell^64^ and further analyzed using Morphometrics^65^. Cell length was calculated from the resulting contours using the MicrobeTracker meshing algorithm^66^. Fractional length contraction during each hyperosmotic shock was calculated from cell lengths immediately before and during the shock and used as a measure of cell-envelope stiffness, with greater contraction indicating lower stiffness.

### Gene expression analysis by RT-qPCR

Transcript abundance of *marA*, *acrB*, and *tolC* was quantified by reverse transcription followed by quantitative PCR (RT-qPCR) according to MIQE guidelines^67^. Cultures were grown in 50-mL glass flasks under standard conditions to mid-exponential phase (OD_600_∼0.2–0.6). Cells were harvested by centrifugation, immediately resuspended in NucleoProtect RNA Reagent (Machery-Nagel), and stored at –80 °C until processing. Cells were disrupted using a FastPrep®-24 Classic bead-beating system (MP Biomedicals).

Total RNA was isolated using a PureLink RNA Mini Kit (Invitrogen) according to the manufacturer’s protocol, including an on-column digestion with RNase-free DNase I (Thermo Fisher Scientific). RNA concentration and purity were assessed using a NanoDrop spectrophotometer, and RNA integrity was assessed by agarose gel electrophoresis based on intact 16S/23S rRNA bands. RNA was stored at –80 °C.

RNA preparations were normalized to 50 ng/μL, and reverse transcription was performed using a High-Capacity cDNA Reverse Transcription Kit (Applied Biosystems). Resulting cDNA was diluted 1:100 in nuclease-free water, and qPCR was performed using iTaq Universal SYBR Green Supermix (Bio-Rad) on a CFX96 Touch Real-Time PCR Detection System (Bio-Rad). Each experiment included no-template and no-reverse-transcription controls. Primer sequences are listed in Table S3. Thermal cycling consisted of 30 s at 95 °C followed by 40 cycles of 5 s at 95°C and 30 s at 60°C. Melt-curve analysis confirmed single-product amplification. *gyrA* and *rpsL* were used as reference genes, and relative transcript abundance was calculated using the 2^−ΔΔ*C*^*^t^* method^68^ following geometric normalization to the reference genes.

## Data Availability

Raw sequencing reads (fastq) for the whole genome sequencing experiments have been deposited into the Sequence Read Archive (SRA) under BioProject accession PRJNA1544651.

## Acknowledgements

The work was supported by NIH R35 GM127331 to P.A.L., NIH F31 GM146401 to R.A.V., an Arnold and Mabel Beckman Foundation Beckman Scholars Award to N.Z., and NSF EF-2125383 to K.C.H. K.C.H. is a Chan Zuckerberg Biohub Investigator. We thank Michael Caparon, Jennifer Philips, Ram Dixit, David Rosen, and Enrique Rojas for discussions, feedback, and suggestions that informed this work. We thank the following people for graciously gifting us strains, plasmids, and alleles used in this study: David Rosen (for *K. pneumoniae* TOP52), Joe Lutkenhaus [for *ftsW*(M269I), *ftsW*(E289G), and the parental S3], Waldemar Vollmer (for *ldtA*::*frt ldtB*::*frt ldtC*::*frt ldtD*::*frt ldtE*::*frt ldtF*::*frt*), Matthew Jorgenson (for pDSW204, pDSW204::MurA, and pDSW204::UppS), Angela Mitchell (for *waaL*::*frt*, *wecA*::*frt*, and *wzzE*::*frt*), and Suckjoon Jun (for the parental strain used to construct the tCRISPRi strains).

## Supplementary Figures

**Figure S1:**
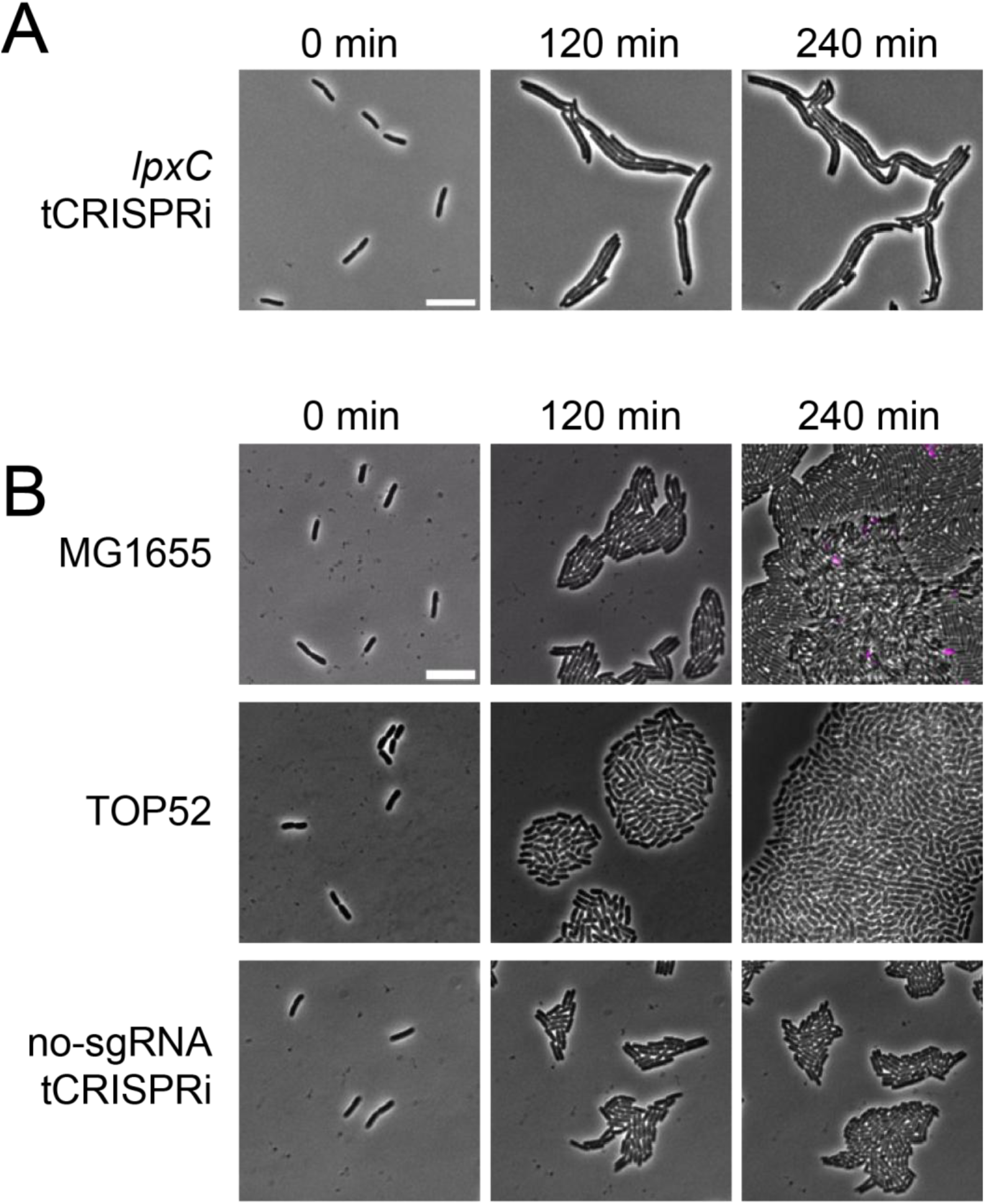
Additional representative images of cells during LPS depletion and control conditions. **(A)** Representative images of a less common morphological phenotype observed following induction of *lpxC* tCRISPRi under standard-osmolarity conditions. Images are representative of two of five independent replicates, with the other three replicates being represented by Fig 2A. Cells exhibit little to no PI staining and form elongated filaments or chains of cells, indicating that LPS depletion can produce substantial morphological changes without detectable loss of membrane barrier integrity. Scale bar, 10 µm. **(B)** Representative images of untreated *E. coli* MG1655 and *K. pneumoniae* TOP52 cells and induced no sgRNA tCRISPRi control cells. Cells proliferate to form confluent microcolonies and exhibit little to no PI staining (magenta), indicating that the morphological abnormalities and loss of envelope integrity associated with LPS depletion are not observed under these control conditions. Scale bar, 10 µm.

**Figure S2:**
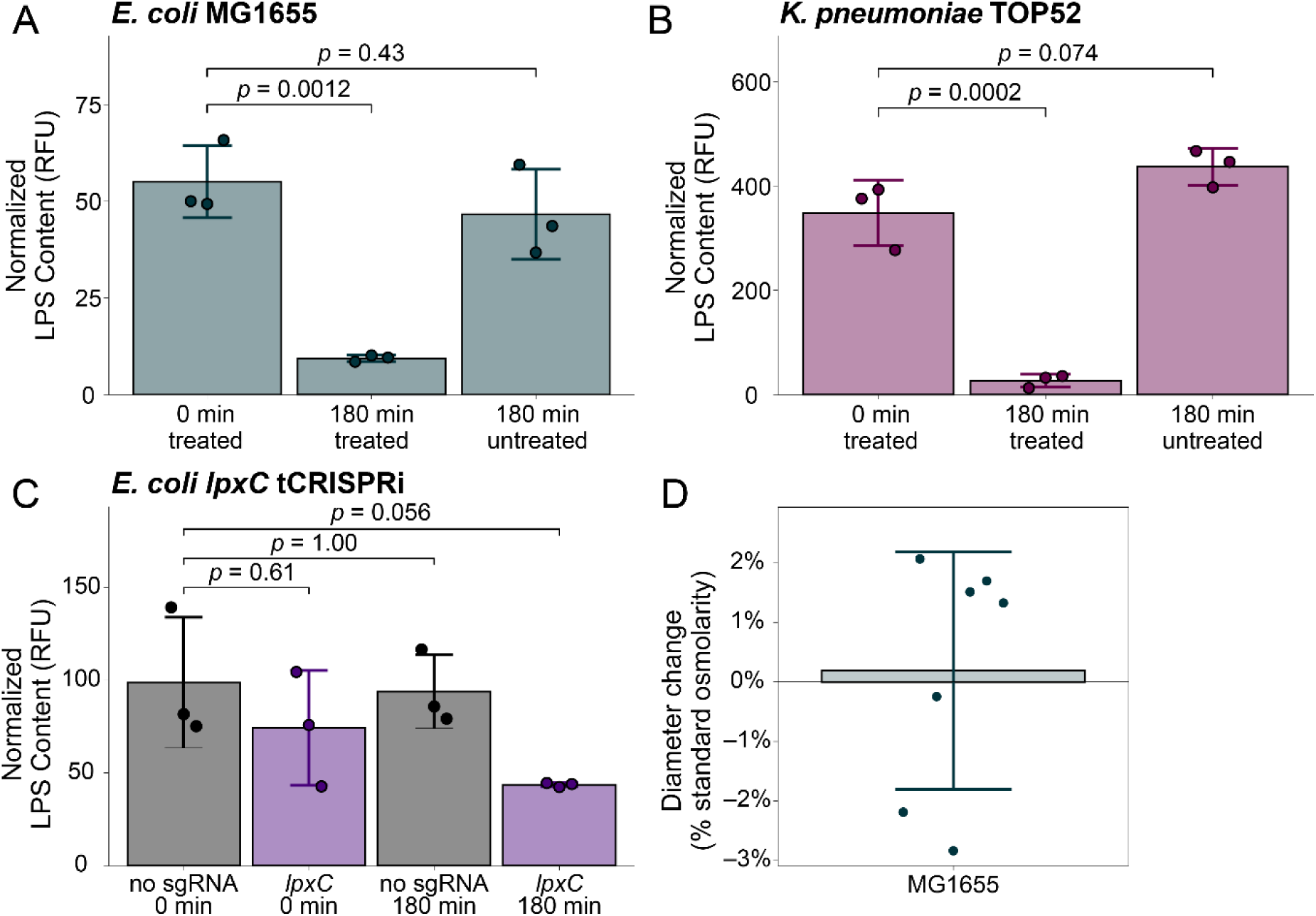
High osmolarity does not prevent LPS depletion or substantially alter CHIR-090 susceptibility. **(A-C)** Quantification of relative LPS abundance during LPS depletion in *E. coli* MG1655 (A), *K. pneumoniae* TOP52 (B), and the *lpxC* tCRISPRi strain (C) grown under high-osmolarity conditions. Values are normalized to a staining control (42-kDa band of the glycoprotein ladder) and total protein content and then scaled such that the mean pretreatment LPS abundance of *E. coli* MG1655 under standard-osmolarity conditions is 100. Each point represents an independent experimental replicate; bars and error bars indicate mean ± SD. *p-*values were determined using Dunnett’s multiple-comparisons test. **(D)** Percent change in CHIR-090 ZOI diameter for MG1655 grown on high-osmolarity medium relative to MG1655 grown under standard-osmolarity conditions. ZOIs were generated using filter-paper disks containing CHIR-090. High osmolarity did not substantially alter CHIR-090 susceptibility (*p-*value = 0.87, determined by Welch’s unequal-variances *t*-test), indicating that its protective effect on envelope integrity is unlikely to result from reduced susceptibility to the inhibitor.

**Table S1:**
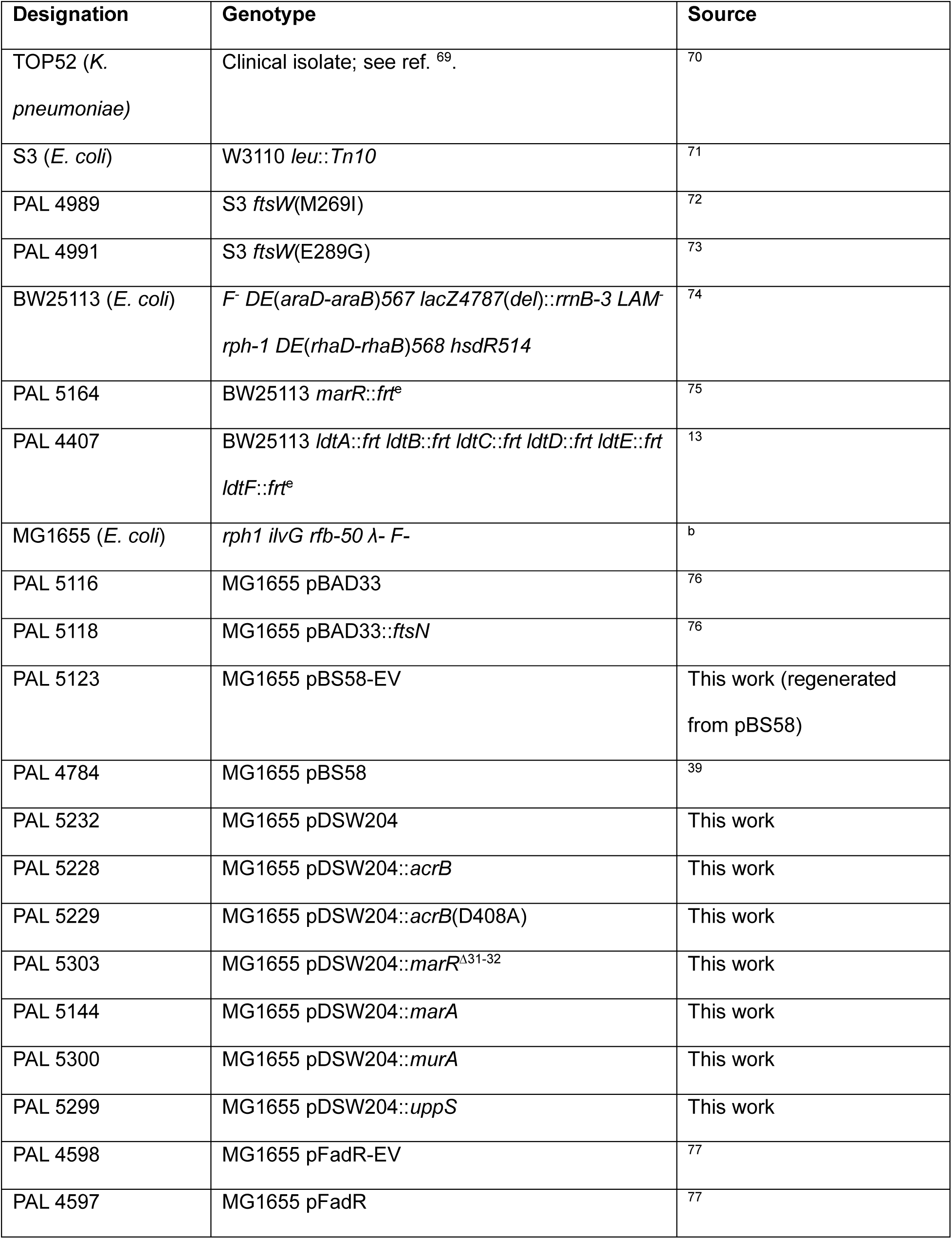
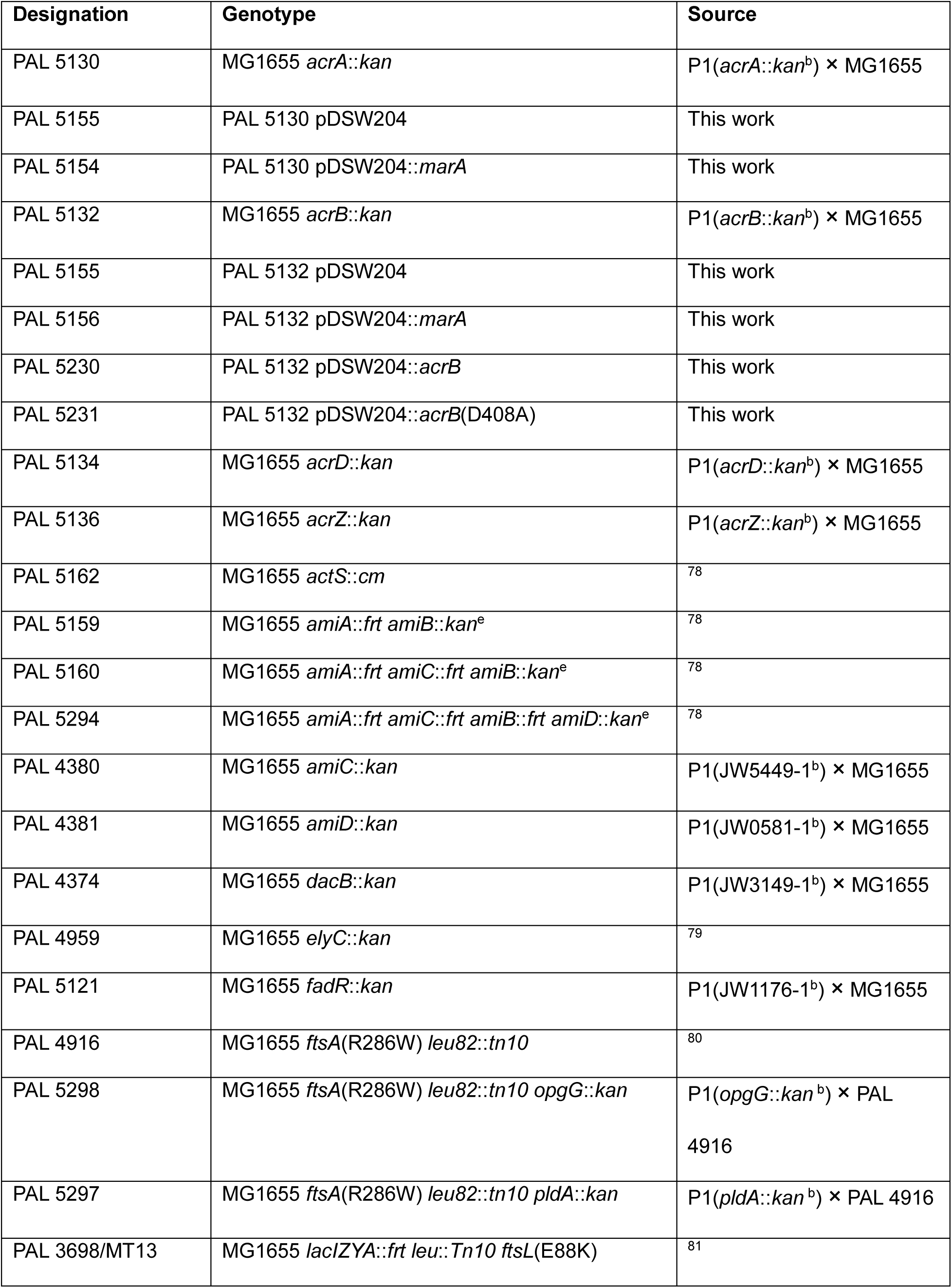
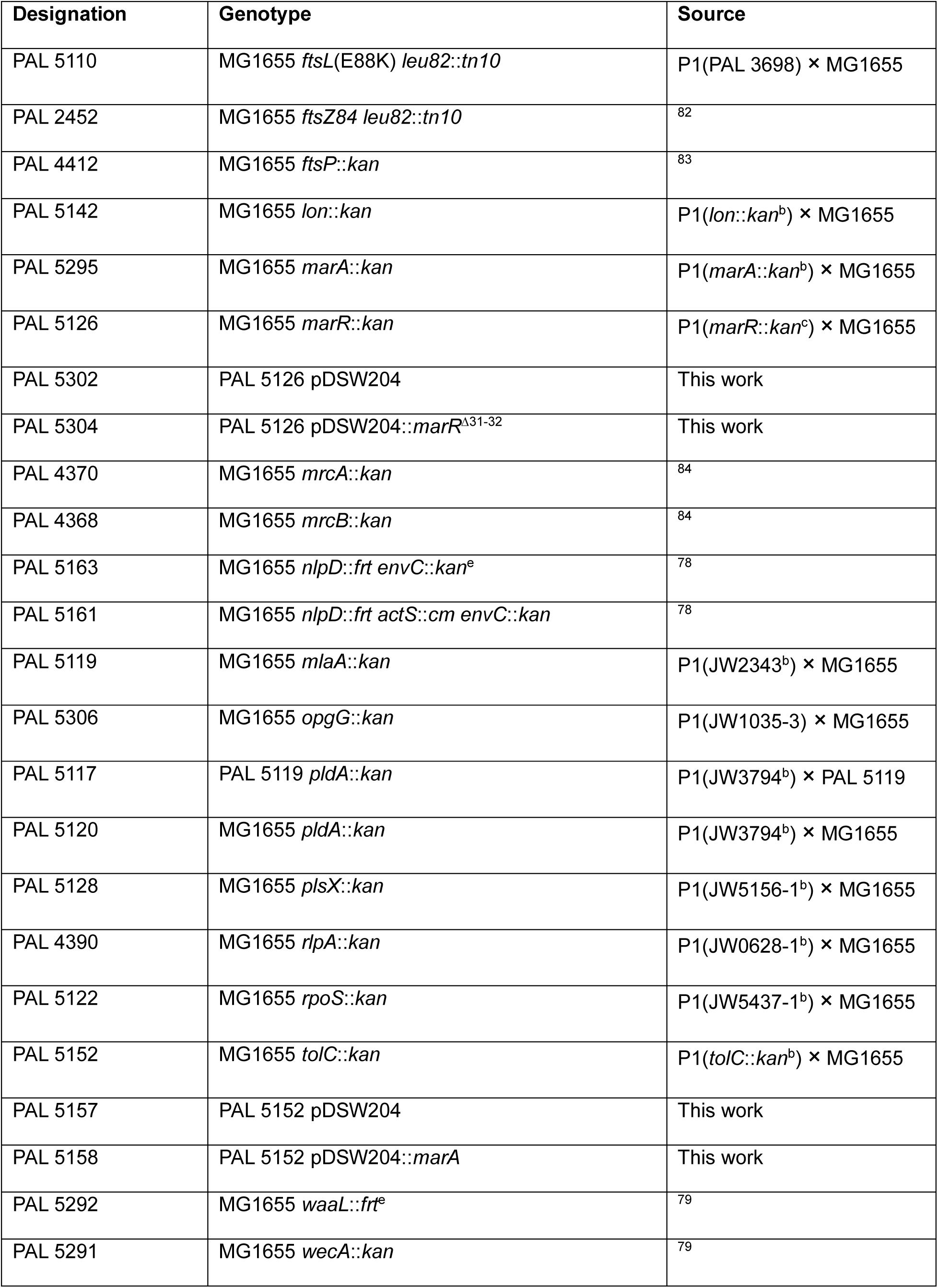
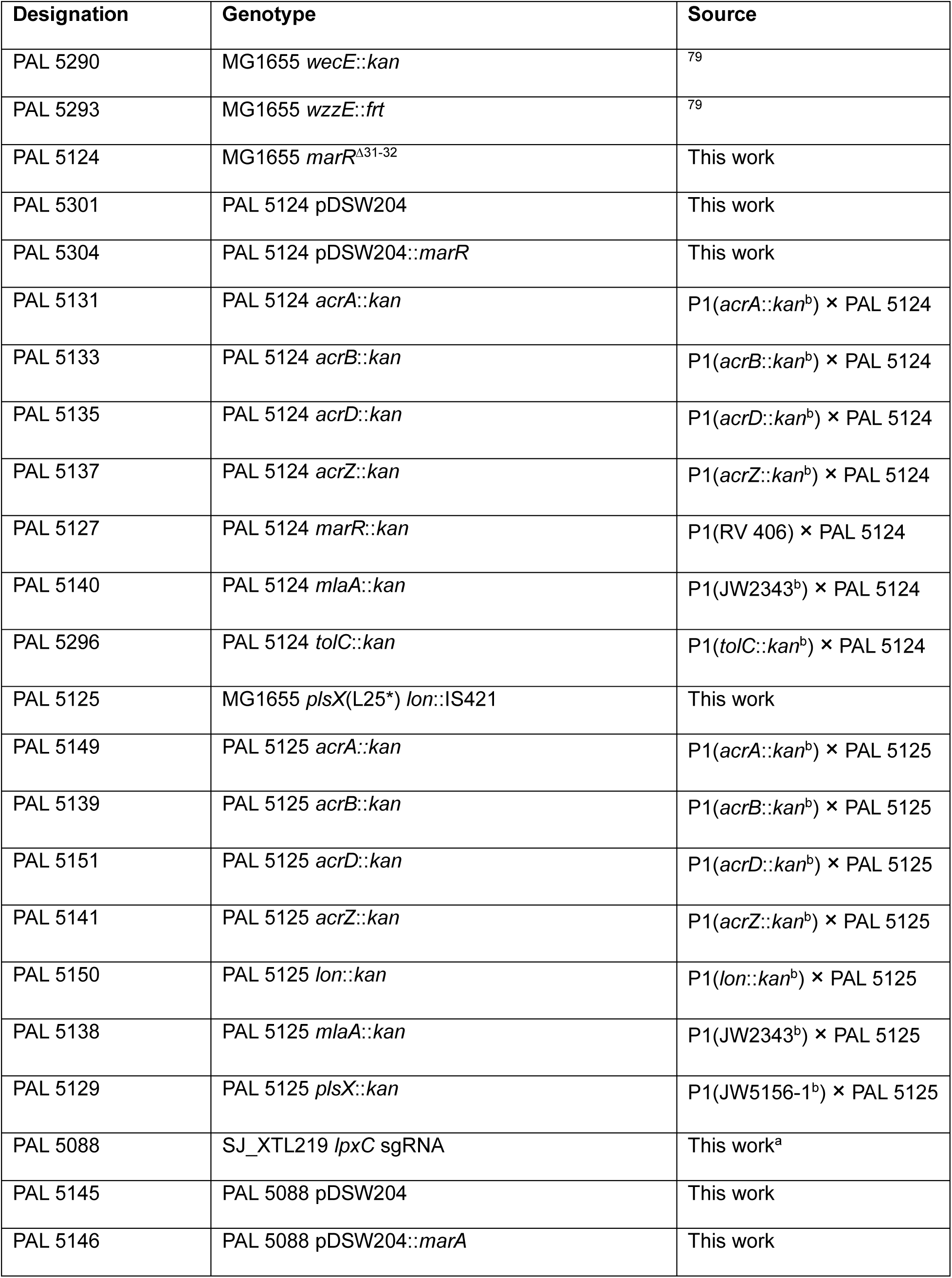
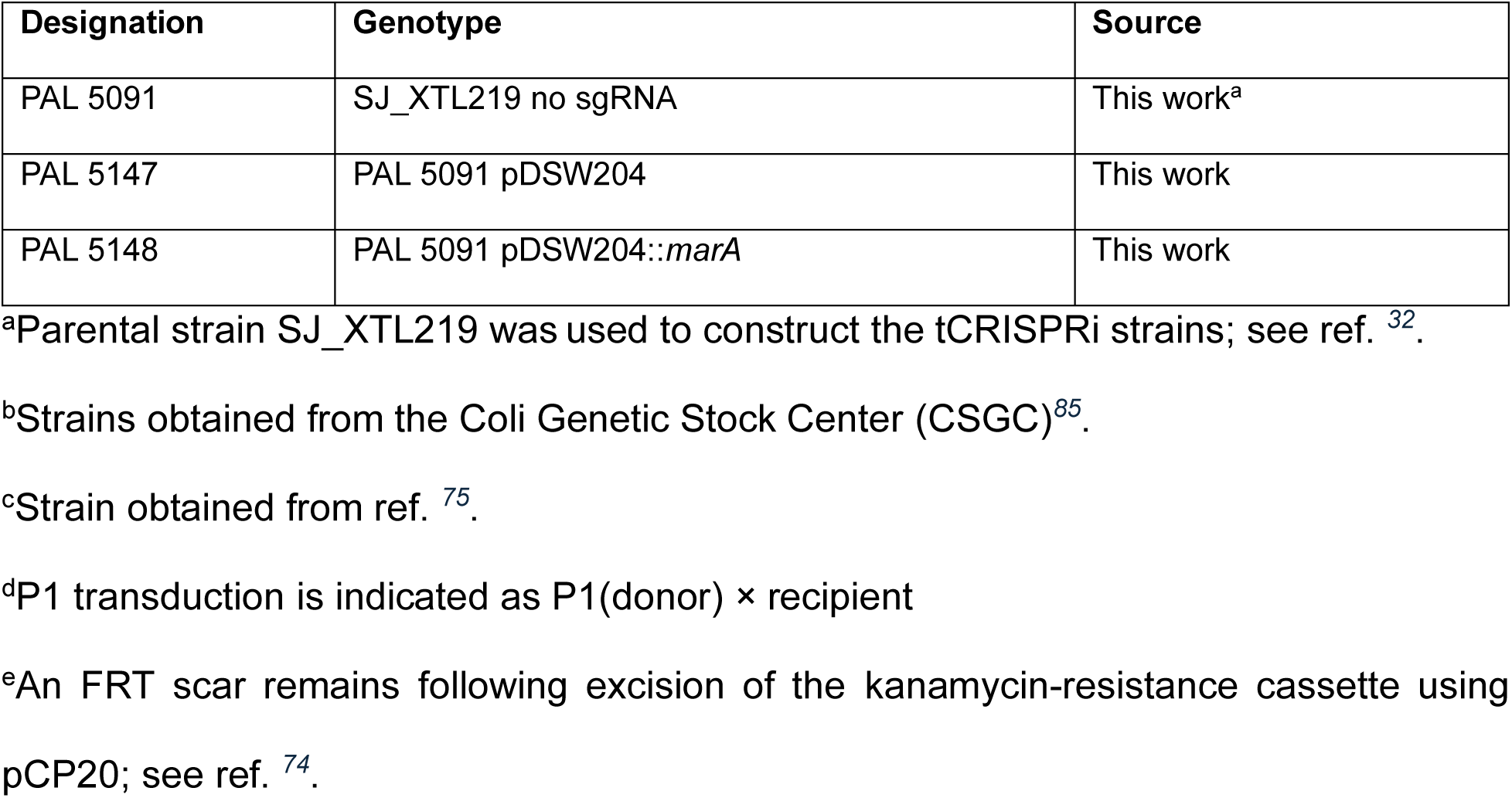
Bacterial strains used in this study.

**Table S2:** Plasmids used in this study.

| Designation | Key Features | Induction conditions | Antibiotic | Source |
| --- | --- | --- | --- | --- |
| pDSW204 (pEV, vector) | pBR ori; *amp<sup>R</sup>*;<br>IPTG regulation<br>(*P<sub>204</sub>*); *lacI<sup>q</sup>* | 200 µM IPTG | 100 µg/mL Amp | <sup>86</sup> |
| pDSW204::*acrB* (*pacrB*) | pDSW204<br>*P<sub>204</sub>::acrB* | 200 µM IPTG | 100 µg/mL mp | This work |
| pDSW204::*acrB*(D408A)<br>[*pacrB*(D408A)] | pDSW204<br>*P<sub>204</sub>::acrB*(D408A) | 200 µM IPTG | 100 µg/mL Amp | This work |
| pDSW204::*marA* (*pmarA*) | pDSW204 *P<sub>204</sub>::marA* | 200 µM IPTG | 100 µg/mL Amp | This work |
| pDSW204::*marR* (*pmarR*) | pDSW204<br>*P<sub>204</sub>::marR* | none | 100 µg/mL Amp | This work |
| pDSW204::*marR*<sup>Δ31-32</sup><br>(*pmarR*<sup>Δ31-32</sup>) | pDSW204<br>*P<sub>204</sub>::marR*<sup>Δ31-32</sup> | 200 µM IPTG | 100 µg/mL Amp | This work |
| pDSW204::*murA* (*pmurA*) | pDSW204<br>*P<sub>204</sub>::murA* | 200 µM IPTG | 100 µg/mL Amp | <sup>87</sup> |
| pDSW204::*uppS* (*puppS*) | pDSW204<br>*P<sub>204</sub>::uppS* | 200 µM IPTG | 100 µg/mL Amp | <sup>87</sup> |
| pBAD33 | P15A ori; *cm<sup>R</sup>*; L-arabinose<br>regulation<br>(*P<sub>araBAD</sub>*); *araC* | 0.2% L-arabinose | 35 µg/mL Cm | <sup>76</sup> |
| pBAD33::*ftsN* (*pftsN*) | pBAD33<br>*P<sub>araBAD</sub>::ftsN* | 0.2% L-arabinose | 35 µg/mL Cm | <sup>76</sup> |
| pBS58-EV (*pftsQAZ*-EV) | pGB2 ori; *aadA* | NA | 100 µg/mL Spec | This work<br>(regenerated<br>from pBS58) |
| pBS58 (*pftsQAZ*) | pBS58-EV *ftsQAZ* | NA | 100 µg/mL Spec | <sup>39</sup> |
| pFadR-EV (*pfadR*-EV) | colE1 ori; *kan<sup>R</sup>*;<br>IPTG regulation<br>(*P<sub>lac</sub>*) | 1 mM IPTG | 25 µg/mL Kan | <sup>77</sup> |
| pFadR (*pfadR*) | pFadR-EV<br>*P<sub>lac</sub>::fadR* | 1 mM IPTG | 25 µg/mL Kan | <sup>77</sup> |

Amp, ampicillin; Cm, chloramphenicol; Spec, spectinomycin; Kan, kanamycin.
Alternative plasmid designations used throughout the manuscript are indicated in
parentheses.

**Table S3:**
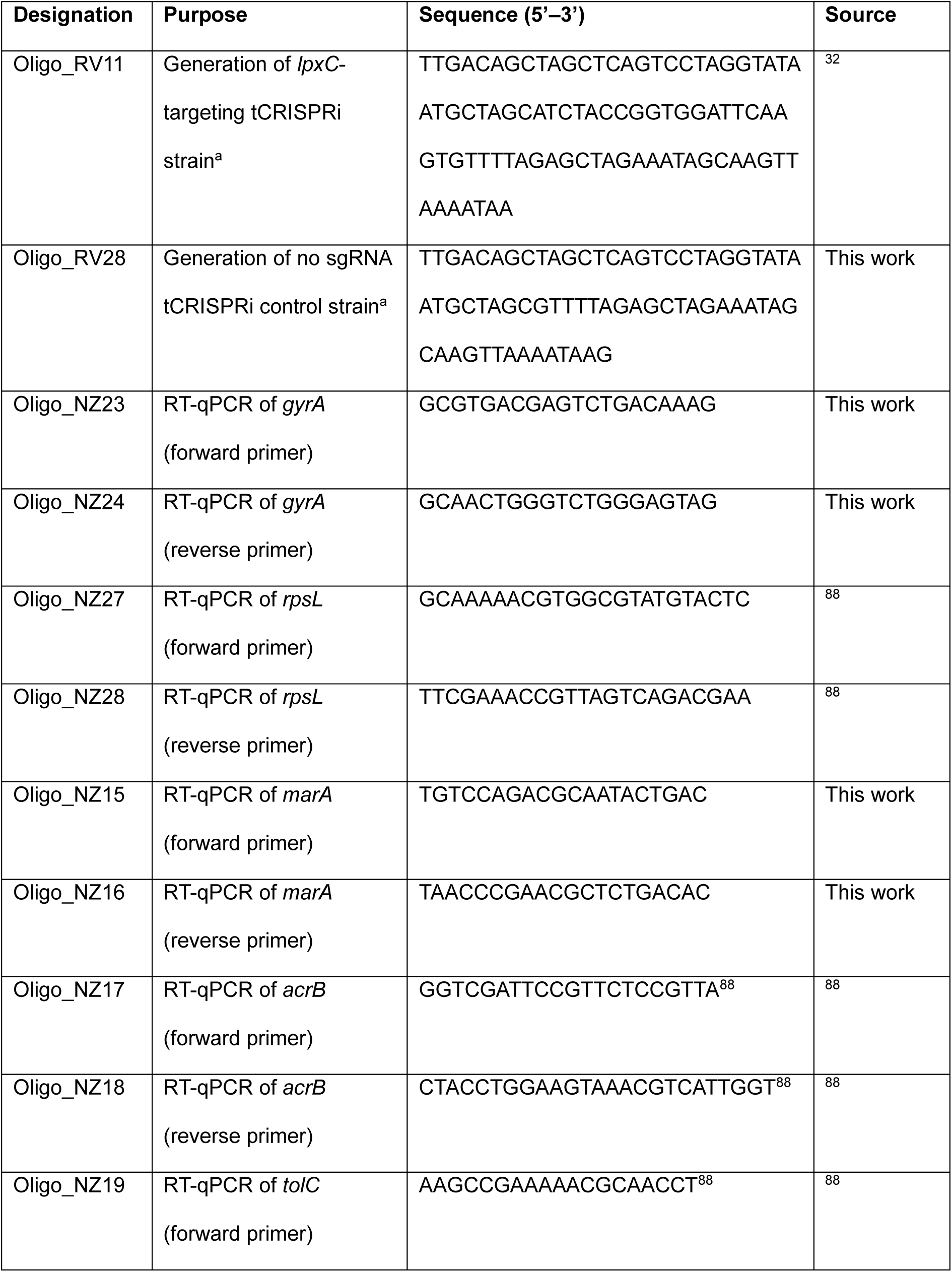
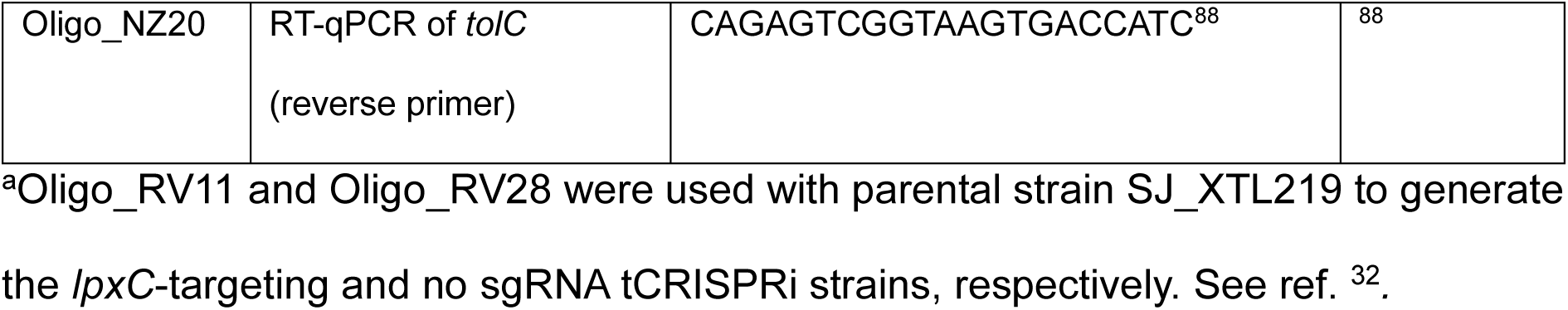
Oligonucleotides used in this study.

**Supplemental Table S4:**
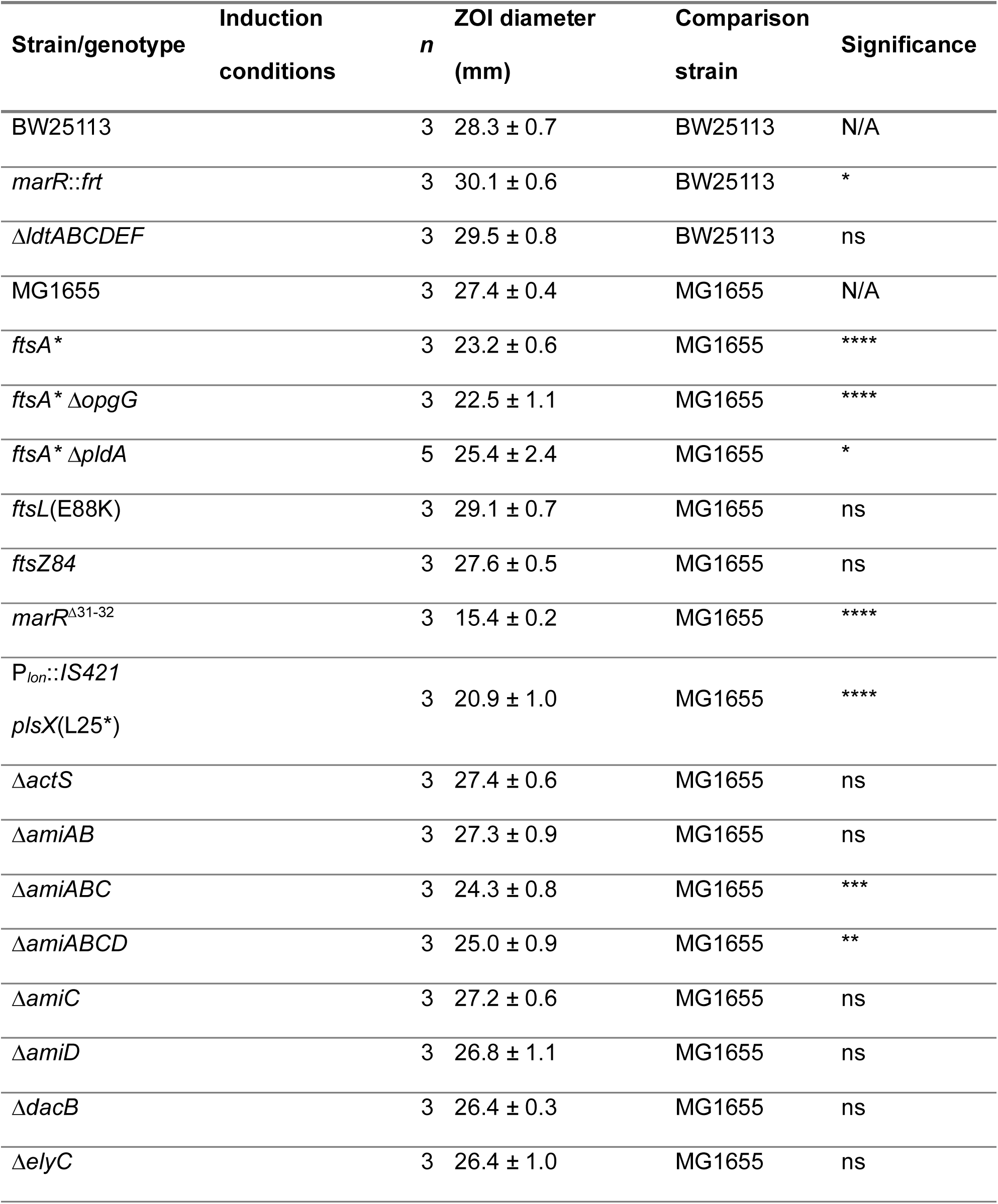
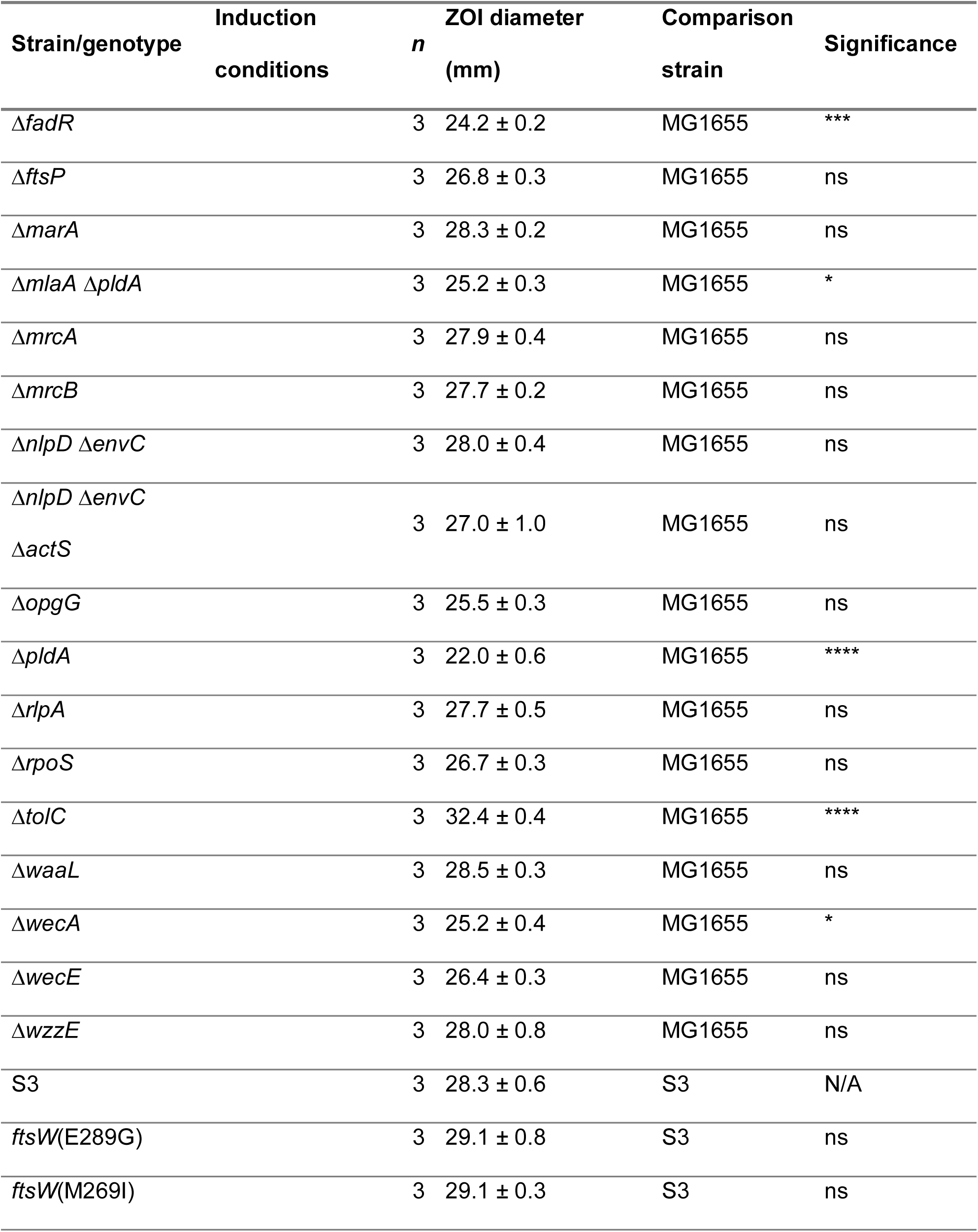
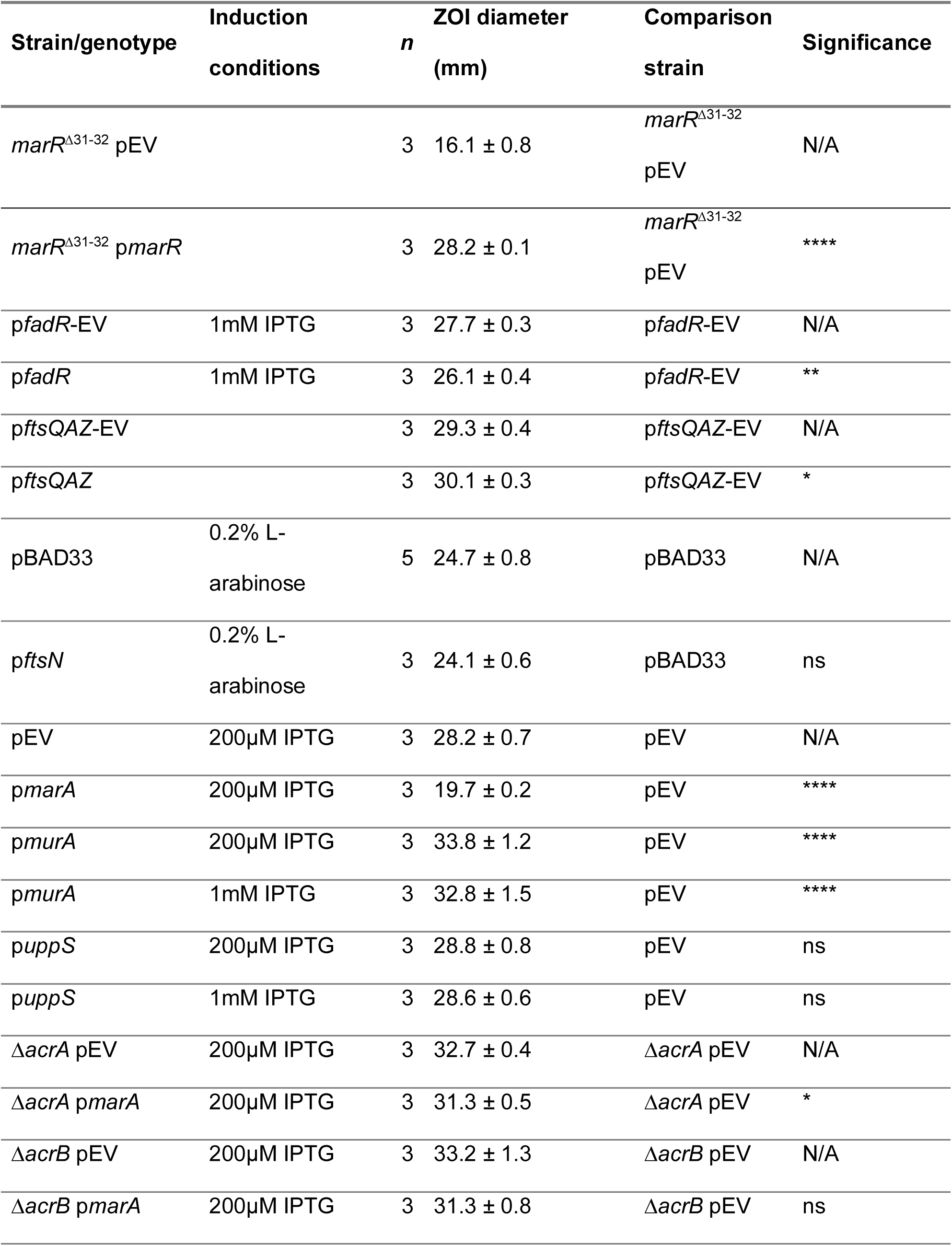
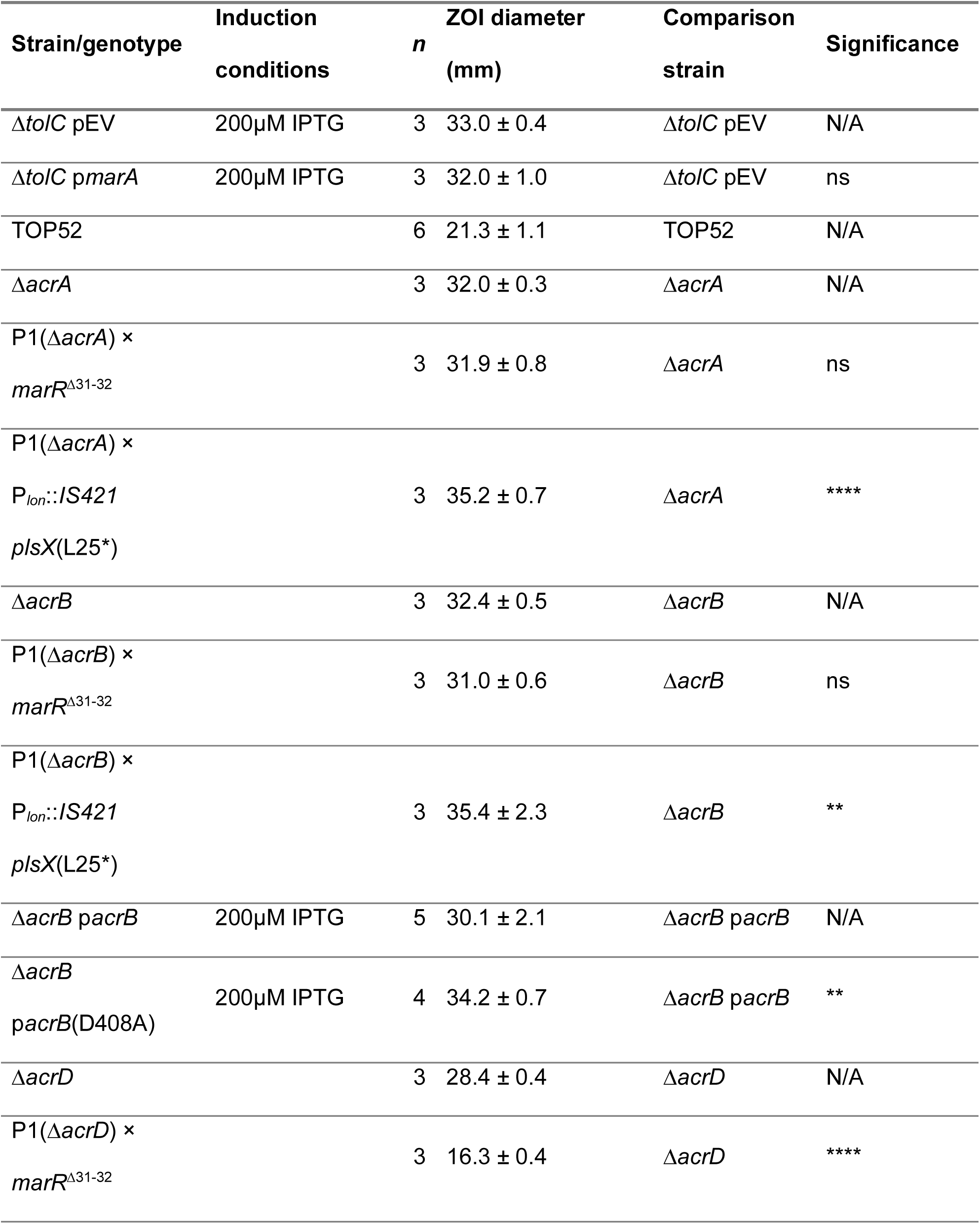
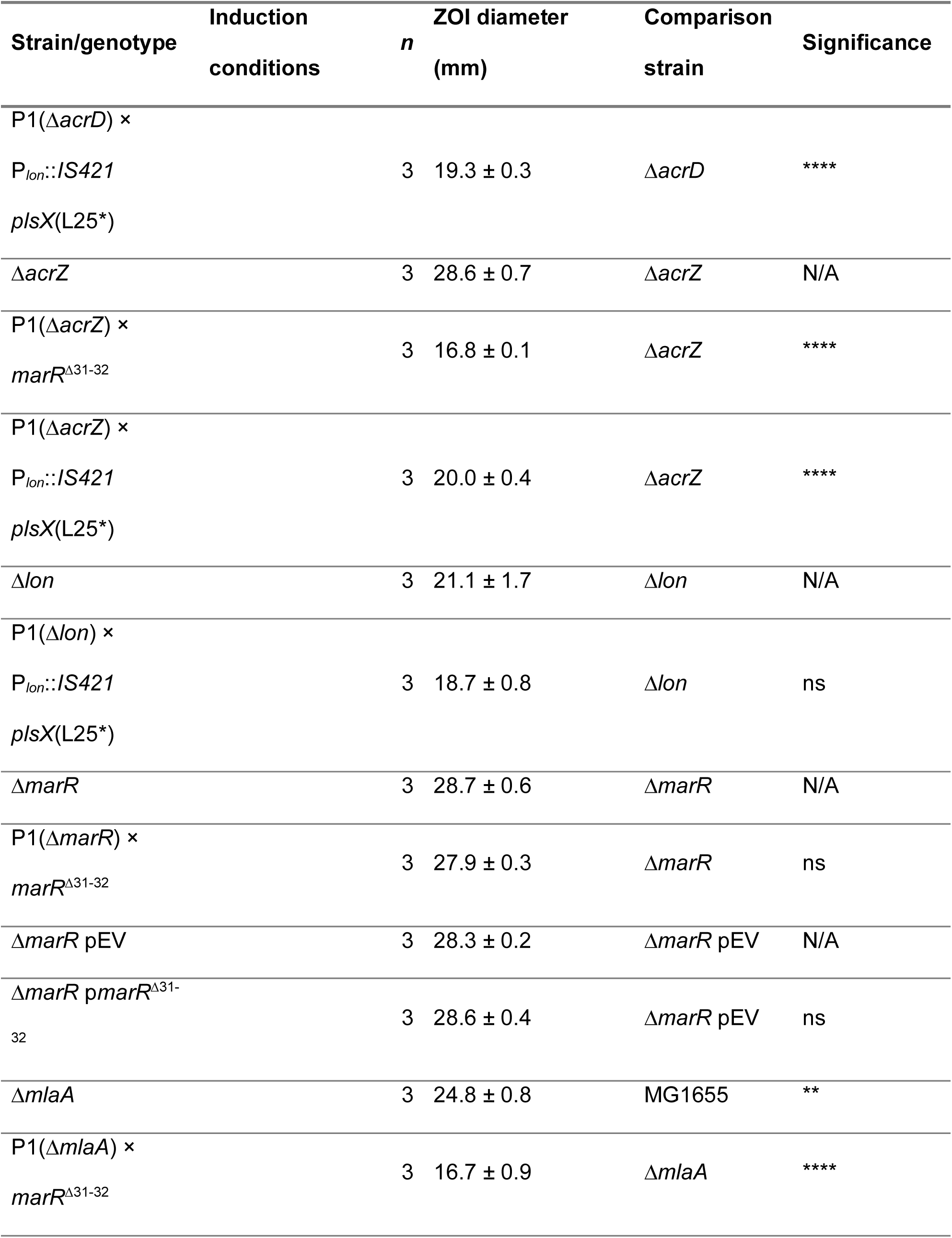
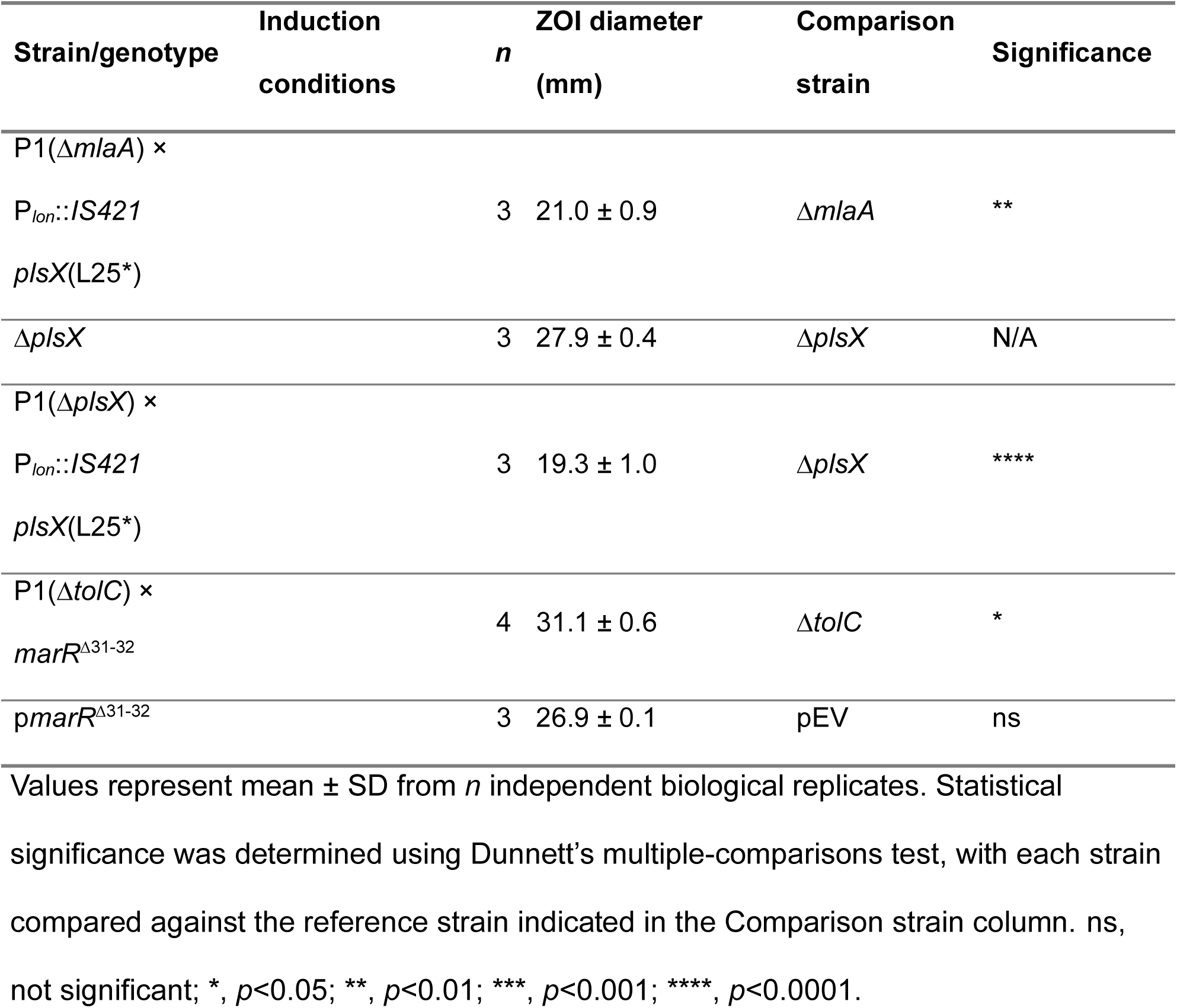
CHIR-090 zones of inhibition (ZOI) across genetic backgrounds and induction conditions.

**Table S5:** Minimum inhibitory concentrations of CHIR-090 under standard and high-osmolarity conditions.

| Strain | LB MIC ($\mu\text{g/mL}$) | LB + 500 mM D-sorbitol MIC ($\mu\text{g/mL}$) |
| --- | --- | --- |
| *E. coli* MG1655 | $0.086 \pm 0.032$ ($n=8$) | $0.088 \pm 0.034$ ($n=5$) |
| *K. pneumoniae* TOP52 | $0.38 \pm 0.14$ ($n=4$) | $0.21 \pm 0.07$ ($n=3$) |

Values represent as mean ± SD from $n$ independent biological replicates.

